# Self-Supervised Natural Image Reconstruction and Large-Scale Semantic Classification from Brain Activity

**DOI:** 10.1101/2020.09.06.284794

**Authors:** Guy Gaziv, Roman Beliy, Niv Granot, Assaf Hoogi, Francesca Strappini, Tal Golan, Michal Irani

## Abstract

Reconstructing natural images and decoding their semantic category from fMRI brain recordings is challenging. Acquiring sufficient pairs of images and their corresponding fMRI responses, which span the huge space of natural images, is prohibitive. We present a novel *self-supervised* approach that goes well beyond the scarce paired data, for achieving both: (i) state-of-the art fMRI-to-image reconstruction, and (ii) first-ever large-scale semantic classification from fMRI responses. By imposing cycle consistency between a pair of deep neural networks (from image-to-fMRI & from fMRI-to-image), we train our image reconstruction network on a large number of “unpaired” natural images (images without fMRI recordings) from many novel semantic categories. This enables to adapt our reconstruction network to a very rich semantic coverage without requiring any explicit semantic supervision. Specifically, we find that combining our self-supervised training with *high-level perceptual losses*, gives rise to new reconstruction & classification capabilities. In particular, this perceptual training enables to classify well fMRIs of never-before-seen semantic classes, *without requiring any class labels during training*. This gives rise to: (i) Unprecedented image-reconstruction from fMRI of never-before-seen images (evaluated by image metrics and human testing), and (ii) Large-scale semantic classification of categories that were never-before-seen during network training. *Such large-scale (1000-way) semantic classification from fMRI recordings has never been demonstrated before*. Finally, we provide evidence for the biological consistency of our learned model.

## 1 Introduction

Natural images span a vastly rich visual and semantic space that humans are experts at processing and recognizing. An intriguing inverse problem is to decode images seen by a person and their semantic categories directly from brain activity (Fig 1a) [1, 2]. Modeling the stimulus as a function of cortical activation (decoding) complements the conventional neuroscientific paradigm of modeling cortical activation as a function of the stimulus (encoding) [1]. It provides us with an additional benchmark of our understanding of the cortical visual code and is thus a cornerstone in cognitive neuroscience [3–10]. Specifically, reconstruction and classification of seen images provide a window into top-down perceptual processes whose probing was previously limited to subjective report. These include visual recall [11], visual imagery in wakefulness [11–15] and sleep [16, 17]. Last, advancements in fMRI-based decoding may be useful for other neuroimaging modalities that provide opportunities for effective brain-machine interfaces [18–20] or for diagnosing disorders of consciousness [21, 22].

**Figure 1.**
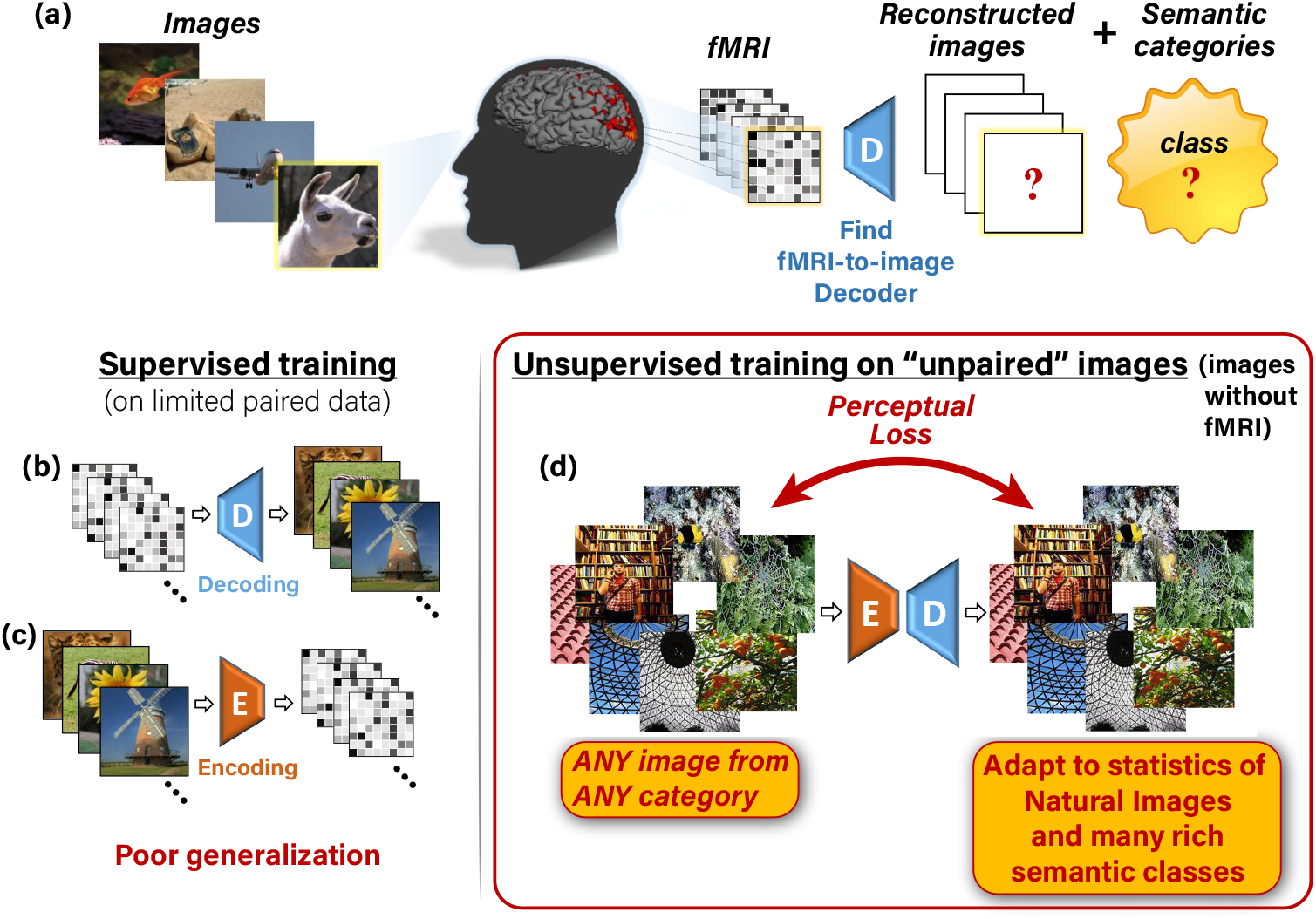
Our self-supervised approach. **(a)** The task: reconstructing images and classifying their semantic category from evoked brain activity, recorded via fMRI. **(b,c)** Supervised training for decoding (**b**) and encoding (**c**) using limited training pairs. This gives rise to poor generalization. **(d)** Illustration of our added self-supervision, which enables training on “unpaired images” (any natural image with no fMRI recording). This self-supervision allows adapting the decoder to the statistics of natural images and many rich semantic classes.

In the image reconstruction task, one attempts to decode natural images which were observed by a human subject from the induced brain activity captured by functional magnetic resonance imaging (fMRI). To learn the mapping between fMRI and image representation, typical fMRI datasets provide many pairs of images and their corresponding fMRI responses, referenced to as “paired” data. The goal is to learn an fMRI-to-image decoder which generalizes well to reconstructing images from novel “test-fMRI”, fMRI response induced by novel images from totally different semantic categories than those in the training data (referred to as “test-images”). Moreover, a complementary challenge to reconstructing the underlying image is also to decode its semantic category. *However, the shortage of “paired” training data limits the generalization power of today’s fMRI decoders*. The number of obtainable image-fMRI pairs is bounded by the limited time a human can spend in an MRI scanner. This also results in a limited number of semantic categories associated with fMRI data. Accordingly, most datasets provide only up to a few thousands of (Image, fMRI) pairs, which are usually obtained from a small number of image categories. Such limited data cannot span the huge space of natural images and their semantic categories, nor the space of their fMRI recordings. Moreover, the poor spatio-temporal resolution of fMRI signals, as well as their low Signal-to-Noise Ratio (SNR), reduce the reliability of the already scarce paired training data [23–25].

Reconstructing natural images from fMRI was approached by a number of methods, which can broadly be classified into three families: (i) Linear regression between fMRI data and handcrafted image-features (e.g., Gabor wavelets) [26–28], (ii) Linear regression between fMRI data and deep (CNN-based) image-features (e.g., using pretrained AlexNet) [12, 29–31], or latent spaces of pretrained generative models [32–35], and (iii) End-to-end Deep Learning [13, 36–38]. To our best knowledge, methods [12] and [36] are the current state-of-the-art in this field. All these methods inherently rely on the available “paired” data to train their decoder (pairs of images and their corresponding fMRI responses). When trained on limited data, purely supervised models are prone to overfitting, which leads to poor generalization to new test-data (fMRI response evoked by new images). To overcome this problem, recent methods [12, 32–34, 36–38] introduced natural-image priors in the form of Generative Adversarial Networks (GANs) or Variational Auto-Encoders (VAEs). These methods enabled leap advancement in reconstruction quality from fMRI, and tend to produce natural-looking images. Nevertheless, despite their pleasant natural appearance, their reconstructed images are often *unfaithful* to the actual images underlying the test-fMRI (see Fig 8a).

Prior work on semantic classification of fMRI recordings induced by natural-images, can be characterized as two families: (i) Classifying new images from previously seen categories, and (ii)	Classifying new images from novel never-before-seen categories. In the first, the categories to-be-decoded (of the test data) are represented in the training data [6, 39–42]. This widely-explored family is limited to decode only the few and typically coarse classes (e.g., faces, scenes, inanimate objects, and birds), which are represented in the limited “paired” data used for decoder training. The second family, which was introduced in [43], addresses the much more challenging case, where the test-categories are *novel*, namely, not directly represented in the training data. Under this setting, decoding novel, diverse, and fine-grained semantic categories (e.g., ImageNet [44]) remains a difficult task because of the narrow semantic coverage spanned by the limited paired training data.

To cope with this data limitation, recent approaches [29, 39, 41–43] harnessed a *pretrained* and semantically separable embedding. In this approach, voxel responses are linearly mapped to *higher-level* feature representation of an image classification network. Once mapped, categorization is achieved by either (i) forward-propagating the decoded representation to the classification layer [29], or by (ii) nearest-neighbor classification against a gallery of category representatives, which are the mean feature representations of many natural images from that category [43]. While these methods benefited from the wide-coverage of their semantic representation (which stems from pretrained image features independently of the fMRI data), their decoding method remained **supervised** in essence. This is because their training of the mapping from fMRI-to-feature representation relies solely on the limited “paired” training data. Consequently, they are prone to poor generalization and are still limited by the poor category coverage of the “paired” training data.

We present a new approach to overcome the limitations mentioned above and inherent lack of training data, simultaneously for both tasks – image reconstruction as well as for *large-scale semantic classification*. We achieve this by introducing *self-supervised training on <u>unpaired images</u> (images without fMRI recordings)*. Our approach is illustrated in Fig 1. We train two types of deep neural networks: an Encoder *E*, to map natural images to their corresponding fMRI response, and a Decoder *D*, to map fMRI recordings to their corresponding images. Composing those two networks, E-D, yields a combined network whose input and output are the same image (Fig 1d). ***This allows for unsupervised training on <u>unpaired images</u>*** (i.e., images without fMRI recordings, e.g., 50,000 natural images from 1000 semantic categories in our experiments). Such self-supervision adapts the Decoder to the statistics of novel natural images and their novel categories (including categories not represented in the “paired” training data or even in the “unpaired” natural images). Importantly, we combine our self-supervised approach with semantics learning ***without requiring any explicit class labels in the training***. This is achieved via: (i) Using perceptual similarity reconstruction criteria. These criteria require for the reconstructed image to be similar to the original image not only at a pixel level, but also that the two images be ***perceptually similar***, in their higher-level deep “semantic” representations. (ii) Designing an Encoder architecture that benefits from a higher-level “semantic” representation of a backbone network that was trained for image classification. Fig 2 exemplifies the power of adding unsupervised training on unpaired images together with the perceptual criteria. Furthermore, beyond the dramatic improvement in the image reconstruction task, we show that our combined self-supervised perceptual approach gives rise also to new capabilities in semantic category decoding (against 1000 classes), despite the scarce fMRI-based training data.

**Figure 2.**
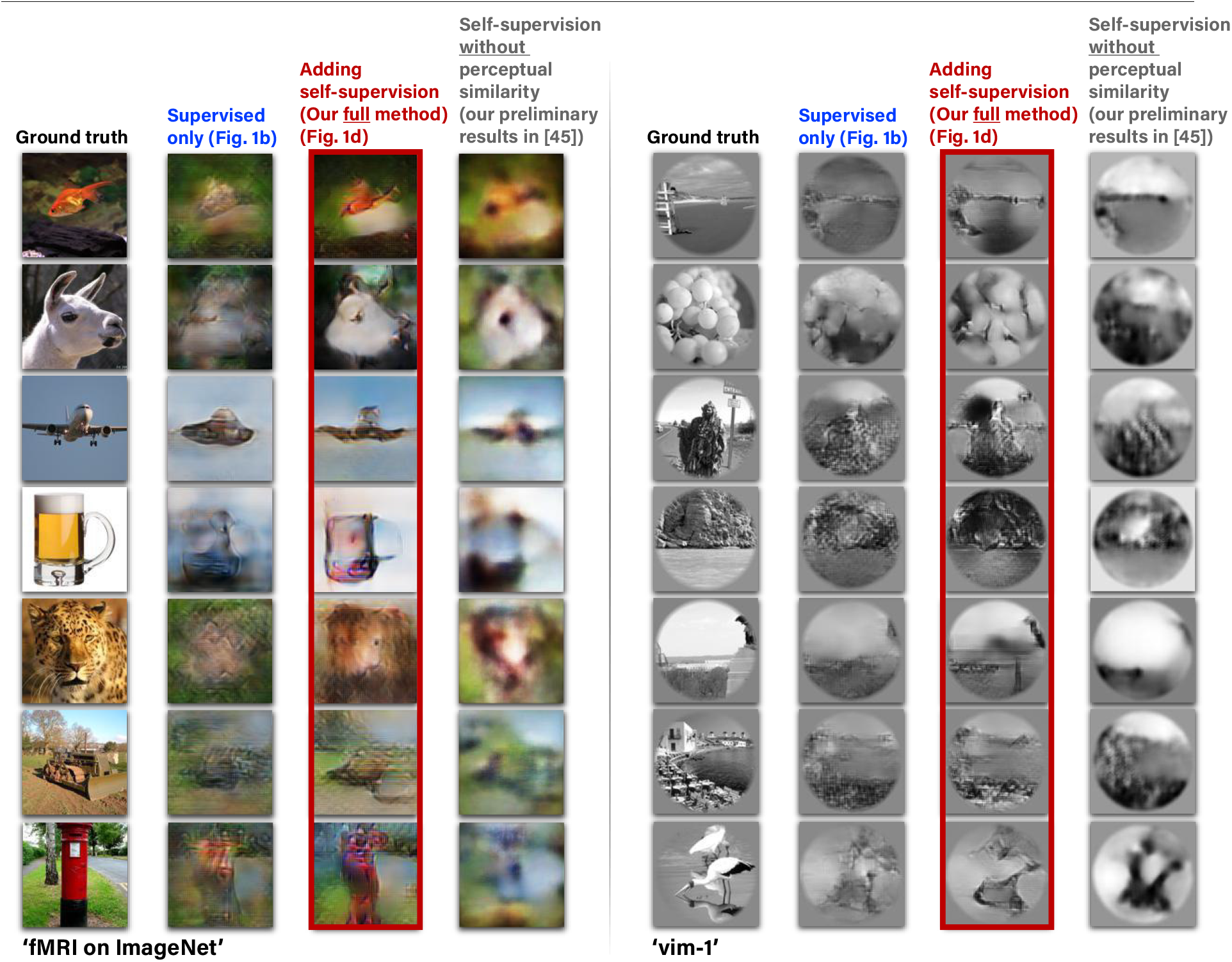
Adding unsupervised training on “unpaired images” together with perceptual criteria improves reconstruction. (Left to Right): • The images presented to the human subjects. • Reconstruction using the training pairs only (Fig 1b). • Reconstruction when adding self-supervised training on unpaired natural images (Fig 1d), and also adding high-level perceptual criteria to the decoder and other important improvements. • Our preliminary results [45] without using the perceptual criteria and other important improvements presented here. Example results are shown for two fMRI datasets: ‘fMRI on ImageNet’ [43] and ‘vim-1’ [26].

Our self-supervision leverages the cycle-consistency nature of our two networks when combined. Notably, our self-supervised training on unpaired natural images differs from an auto-encoder: When training under this configuration the Encoder weights are kept fixed, thus constraining the input images to go through the true fMRI space. Furthermore, unsupervised training on unpaired natural images was also recently proposed in [36], where they used these images to produce additional surrogate fMRI-data to train their model. However, this was never addressed in the context of semantic decoding from fMRI data. Moreover, their image reconstruction criteria were limited to pixel level alone and did not include any type of perceptual or visual-features based reconstruction criteria. To the best of our knowledge, we are the first to present classification of semantic categories that are never-before-seen during training <u>at a large-scale</u> (1000-way – i.e., detecting the correct class out of more than 1000 rich classes).

A preliminary version of our self-supervised approach and partial results (in the context of image reconstruction only) were previously presented in a conference proceeding [45]. However, here we present our complete and advanced reconstruction algorithm that has undergone major extensions, enabling a leap improvement in the image reconstruction quality over our previous method [45] (as demonstrated in Fig 2). Moreover, our new extended framework now enables also impressive *semantic classification capabilities* from fMRI data. As an aid for readers familiar with our previous report [45], we list the four major extensions introduced in the present paper: (i) We introduced two significant improvements to our algorithm: Adding high-level perceptual criteria [46] on the reconstructed images (in contrast with optimizing Mean-Square-Error loss, and on low-level features alone), and increasing the expressiveness of the Encoder architecture to include higher-level “semantic” representations by using multiple levels of the pretrained VGG network (in contrast with a single readout layer before). Our present algorithm provides state-of-the-art reconstructions compared to leading existing methods to-date, as evident through image-metric-based comparisons as well as extensive human behavioral evaluations. (ii) We extended the self-supervised approach to allow also for <u>semantic classification</u> of reconstructed images. This classification relies on and demonstrates the fidelity of the reconstructions. (iii) We analyze the contributions of different visual cortex areas to the resulting image reconstructions, and (iv) We evaluate the biological consistency of the learned models.

Importantly, adding the perceptual similarity was powerful enough to shadow and marginalize the previously dominant effect of training on unpaired fMRI data in [45]. Accordingly, training on unpaired fMRI is omitted in the present framework (despite being a legitimate feature enabled by our self-supervised approach).

Our contributions are therefore several-fold:

- A self-supervised approach for simultaneous image reconstruction and semantic category decoding, which can handle the inherent lack of image-fMRI training data.
- Unprecedented state-of-the-art image-reconstruction quality from fMRI of never-before-seen images (from never-before-seen semantic categories).
- Large-scale semantic classification (1000+ rich classes) of *never-before-seen semantic categories*. To the best of our knowledge, such large-scale semantic classification capabilities from fMRI data has never been demonstrated before.
- We provide analyses showing predominance of early visual areas in image reconstruction quality, and demonstrate biologically plausible receptive field formation in our learned models (although such constrains were not imposed in the training, but rather automatically learned by our system).

## 2 Overview of the Approach

This section provides a high-level overview of our approach and its key components. The technical details and the experimental datasets are described in Methods.

### 2.1 Self-supervised image reconstruction from brain activity

The essence of our approach is to enrich the scarce paired image-fMRI training data with easily accessible natural images for which *there are no fMRI recordings*. This type of training is enabled by imposing cycle-consistency on the “unpaired images”, using two networks, which learn two inverse mappings: from images to fMRI (Encoding), and vice versa – from fMRI to images (Decoding).

Our training consists of Encoder training followed by Decoder training, which we define in the two phases illustrated in Fig 3. In the first phase, we apply supervised training of the Encoder *E* alone. We train it to predict the fMRI responses of input images using the image-fMRI training pairs (Fig 3a). In the second phase, we use the pretrained Encoder (from the first phase) and train the Decoder *D*, keeping the weights of *E* fixed (Fig 3b). *D* is trained using both the paired and the unpaired data, simultaneously. Here, each training batch consists of two types of training data: (i) image-fMRI pairs from the training set (Fig 1b), and (ii) unpaired natural images (with no fMRI, Fig 1d).

**Figure 3.**
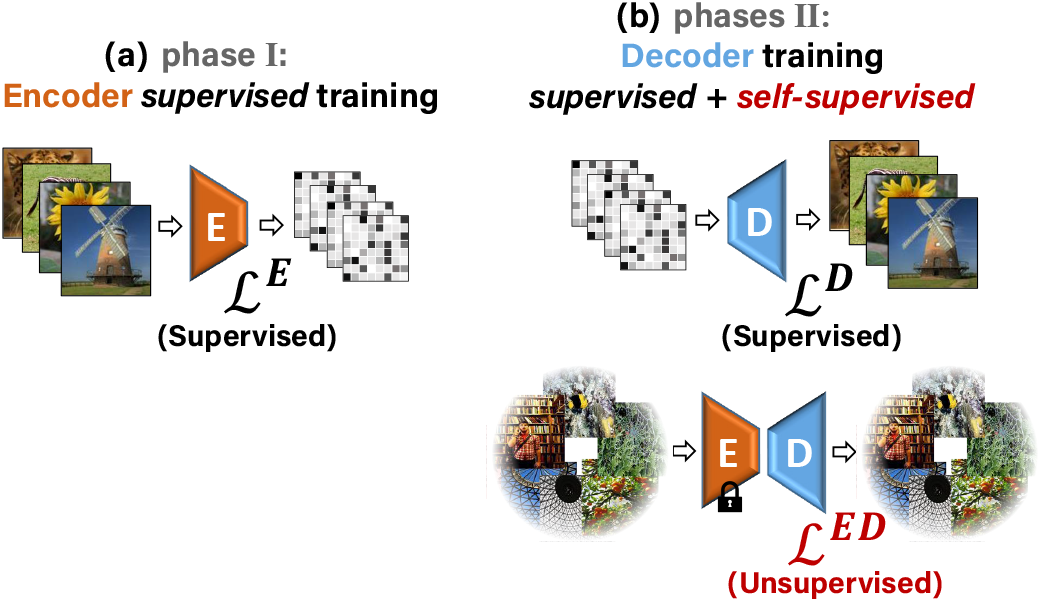
Training phases. **(a)** The first training phase: Supervised training of the Encoder with {Image, fMRI} pairs. **(b)** Second phase: Training the Decoder with two types of data simultaneously: {fMRI, Image} pairs (supervised examples), and unpaired natural images (self-supervision). The pretrained Encoder from the first training phase is kept fixed in the second phase.

Training on *unpaired natural images* (without fMRI) allows to augment the training with data from a much richer semantic space than the one spanned by the paired training data alone. Specifically, we draw the unpaired images from a large *external* database of 49K images from 980 ImageNet (“ILSVRC”) classes, which are mutually exclusive not only to the test-images contained in the fMRI (paired) dataset, but also to their underlying test classes (test categories). In principle, for optimal networks *E* and *D*, the combined *E*-*D* network should yield an output image which is identical to its input image. This should hold for any natural image (regardless if an fMRI was ever recorded for it). Importantly, our image reconstruction losses (in both ℒ_*ED*_ and ℒ_*D*_, Fig 3) require for the reconstructed image to be similar to the original image not only at a pixel level. We further require these two images to be ***perceptually similar***, at a semantic level.

Perceptual Similarity is a metric that highly correlates with *human* image-similarity perception, and involves a broad range of visual feature representation levels [46]. Our Perceptual Similarity loss is illustrated in Fig 4. We apply a pre-trained VGG classification network (a network which was trained for the task of object recognition from images [47]), on the reconstructed and the original images. We then impose similarity between their corresponding deep-image-features, extracted from multiple deep layers of VGG. *Using this metric as our reconstruction criterion enables to learn low-level to high-level “semantic” information from the broad semantic space, which is spanned by the external database of ‘unpaired’ images and their classes; It, thus, encourages the Decoder to output images that are not only accurate at low-level visual features, but are also semantically meaningful.* Importantly, the <u>class labels</u> of the unpaired images (or the paired images) <u>are never used in our training process, hence may be unknown</u>. Adding the perceptual loss in combination with training on additional unpaired images gives rise to a leap improvement in the fMRI-to-image reconstruction quality compared to any previous method (including our previous method [45] – see Fig 2, as well as compared with other methods – see Fig 8). It provides a dramatic improvement in detail level and perceptual interpretability of the reconstructed images. Our new approach further enables large-scale semantic classification of fMRI into rich novel semantic categories.

**Figure 4.**
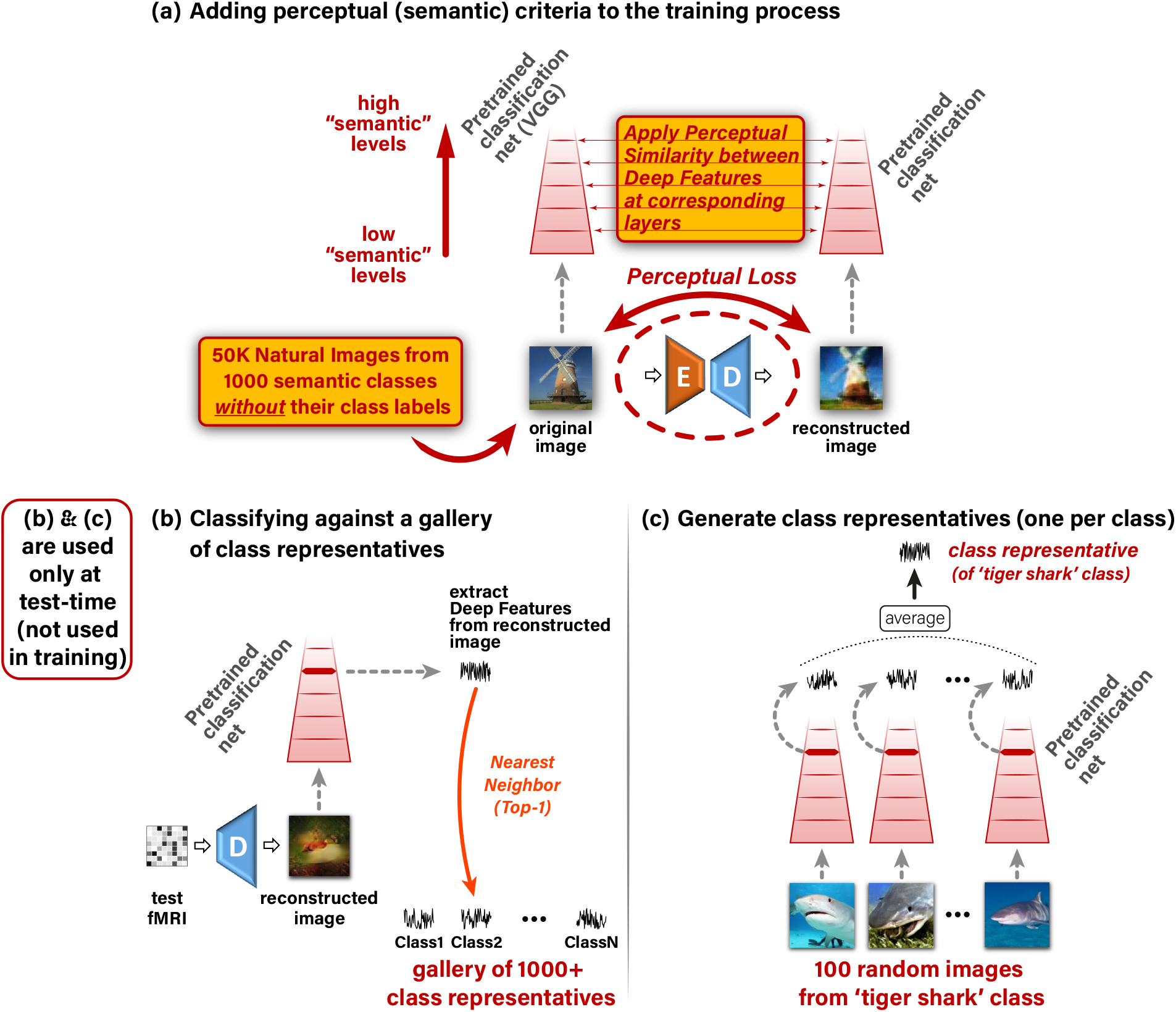
Adding high-level perceptual criteria improves reconstruction accuracy and enables large-scale semantic classification. **(a)** Imposing Perceptual Similarity on the reconstructed image at the Decoder’s output, as applied when training on unpaired natural images (<u>without</u> fMRI and <u>without</u> any class labels) from many novel semantic classes. This adapts the Decoder to a significantly broader semantic space despite not having any explicit semantic supervision. **(b)** To classify a reconstructed image to its novel semantic class we extract Deep Features using a pretrained classification network, and follow a nearest-neighbor class-centroid approach against a large-scale gallery of 1000+ ImageNet classes. **(c)** We define class representatives as the mean-embedding of many same-class images [43].

### 2.2 Self-supervised image classification to semantic categories

Our new self-supervised perceptual approach extends well beyond the task of image reconstruction. It further allows for large-scale semantic classification of fMRI data. We present classification of fMRI data against a gallery of more than 1000 rich image classes, in a challenging 1000-way classification task (see Fig 5). **Such large-scale semantic classification of fMRI data (to 1000-way, with promising results) has never been demonstrated before.**

**Figure 5.**
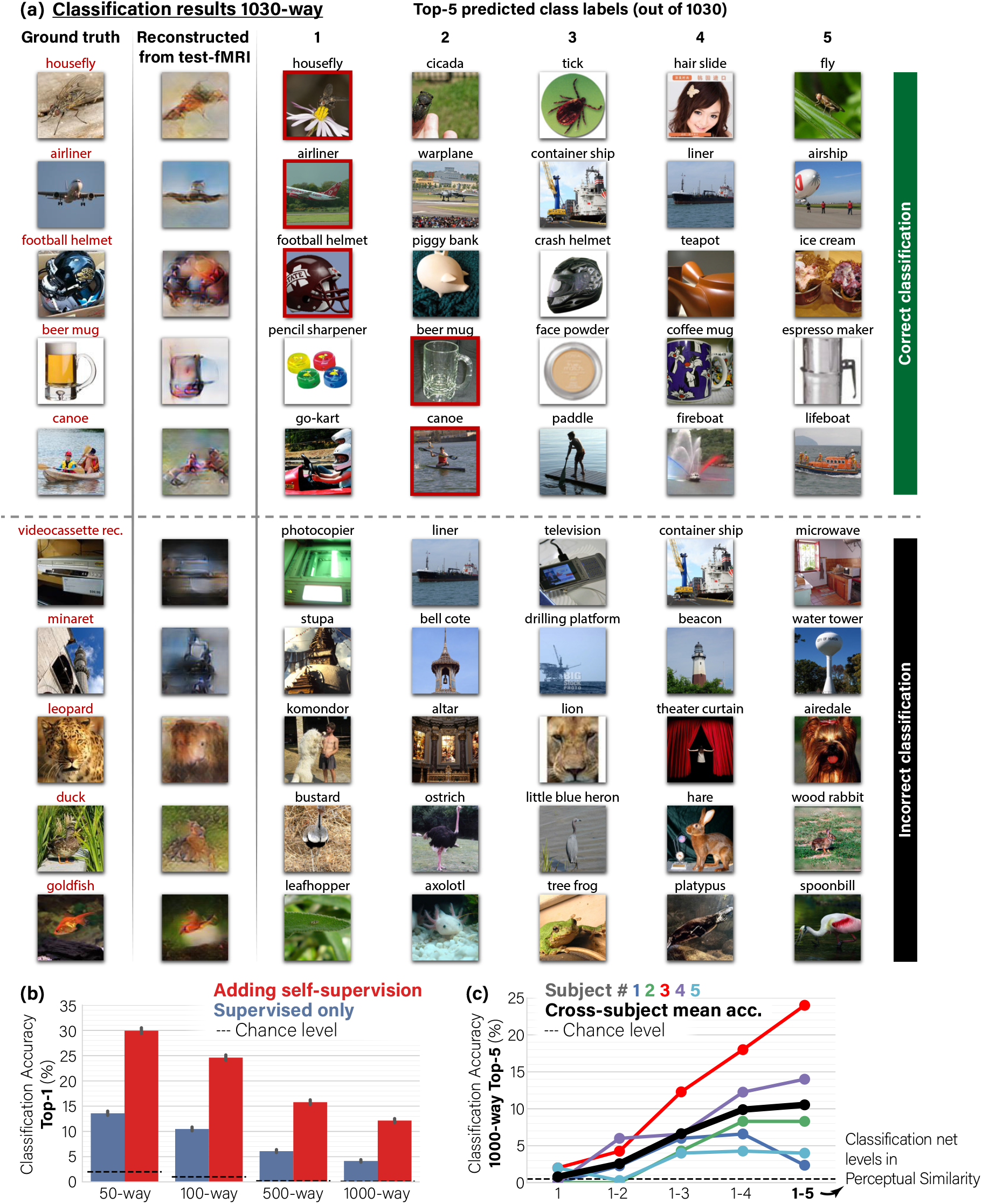
Self-supervision allows classification to rich and <u>novel</u> semantic categories. **(a)** Visual classification results showing the Top-5 predictions out of 1030 classes for reconstructed test-image. We show examples where the ground-truth class (marked in red) is ranked among the Top-5 **(correct classification)** or excluded from it **(incorrect classification)**. For visualization purposes only, each class is represented by the nearest-neighbor image from the 100 randomly sampled images of the particular class. Note that ”incorrect” predicted classes are often reasonable (e.g., ”Leopard” wrongly predicted as ”Lion”; ”Duck” wrongly predicted as ”Ostrich”). **(b)** Top-1 Classification accuracy in an n-way classification task. Adding unsupervised training on unpaired data (Fig 1d,e) dramatically outperforms the baseline of the supervised approach (Fig 1b). **(c)** Ablation study of the Classification accuracy as a function of the Perceptual Similarity criterion for decoder training: Applying “partial” perceptual similarity using only the outputs of the first VGG16 block (low ”semantic” layers), and up to all its 5 blocks (high ”semantic” layers). Applying full Perceptual Similarity on higher-level VGG features substantially improves classification performance. Panels **a,b** show results for subject three. 95% Confidence Intervals shown on charts.

Our classification approach is based on our self-supervised perceptual reconstruction method described above. We use our perceptually trained Decoder to reconstruct the test-images from their test-fMRI. We then classify the reconstructed images against 1000+ rich ImageNet semantic classes. Fig 4b,c shows our classification approach. To classify a reconstructed image to its novel semantic class, we match a “Deep-Feature signature” extracted from the reconstructed image, against “class-representative Deep-Feature signatures” (one per class), in a gallery of 1000+ semantic categories, which also include the 50 *novel* test classes. More specifically: (i) We extract Deep Features from the reconstructed image at an intermediate level of a pretrained classification network (Fig 4b), (ii) Following [43], for each class in the gallery, we compute a single “class representative” using 100 randomly sampled images from that class. The class representative is defined as the average Deep Features (centroid) of those 100 randomly sampled images from that class (see Fig 4c). (iii) We compute the correlation between the Deep Features extracted from the reconstructed image and each of the 1000+ class representatives. These yield 1000+ “semantic similarity” scores (Fig 4b). Ranking the gallery classes according to these similarity scores (for each reconstructed image) provides the basis for semantic classification at any desired ‘Top-X’ accuracy level (Fig 5). Specifically, the classification is marked ‘correct’ when the ground truth category obtains the best similarity score among all 1000+ classes (‘Top-1’), or when it is among the top 5 most similar classes (‘Top-5’). The location of the ground-truth class within the sorted list of 1000+ classes further provides a ”rank score” for evaluating our classification accuracy (see Table 2).

**Table 1.** Summary of fMRI datasets used in analyses. Repeat count refers to the number of fMRI recordings per presented stimulus. Voxel count refers to approximated number of voxels used in analysis.

| fMRI Dataset | $N$ train images ($K$ repeats) | $N$ test images ($K$ repeats) | $N$ voxels |
| --- | --- | --- | --- |
| fMRI on ImageNet [43] | 1200 (1) | 50 (35) | 4500 |
| vim-1 [26] | 1750 (2) | 120 (13) | 8500 |

**Table 2.** Self-supervision allows classification to rich and novel semantic categories. Median rank of the ground truth class among 1030 class representatives (Lower is better). Significant differences between the two methods are marked with asterisks (Mann-Whitney test, N = 50, p < .05). Adding self-supervision leads to significant improvement in classification rank for the four (out of five) subjects with the highest fMRI median noise-ceiling. SD and SE are standard deviation and error, respectively.

| Classification rank (out of 1030) for ‘fMRI on ImageNet’ [43] |  |  |  |
| --- | --- | --- | --- |
|  | fMRI data noise-ceiling<br>median (SD) | Supervised only<br>median rank (SE) | Adding self-supervision<br>median rank (SE) |
| **Subject 1** | 0.56 (0.28) | 136.0 (48.3) | **116.5 (39.0)** |
| **Subject 2** | 0.57 (0.30) | 156.5 (48.1) | **\*85.0 (31.1)** |
| **Subject 3** | 0.73 (0.28) | 118.0 (37.0) | **\*67.0 (32.6)** |
| **Subject 4** | 0.68 (0.30) | 165.0 (49.9) | **\*72.0 (25.5)** |
| **Subject 5** | 0.58 (0.29) | 212.5 (40.0) | **\*71.0 (34.9)** |

This classification procedure greatly benefits from our self-supervised perceptual approach, which enables to train on *additional unpaired images from arbitrarily many novel semantic categories* (Fig 1b). This allows to adapt the Decoder to a much richer (practically unlimited) semantic coverage in a **completely category-free way**, namely without any explicit semantic supervision during training. The key component which induces semantically meaningful decoding is the Perceptual Similarity metric, which involves <u>higher-level</u> ”semantic” criteria. This type of <u>non-specific</u> semantic supervision enables our method to generalize well to new never-before-seen semantic classes – classes which are neither contained in the paired training data, nor in the unpaired external images used at train time.

## 3 Methods

### 3.1 Self-supervised Encoder/Decoder training

The Encoder (E) is trained first on predicting fMRI responses from input images. Once trained, the encoder is fixed, and the Decoder (D) is trained on reconstructing images from fMRI responses. The motivation for training the Encoder and Decoder in separate phases (with a fixed Encoder during Decoder training) is to ensure that the Encoder’s output does not diverge from predicting the fMRI responses by the unsupervised training objectives 1d. Additionally, we start by supervised training of the Encoder in order to allow it to converge at the first phase, and then serve as strong guidance for the more severely ill-posed decoding task [9], which is the focus of the next phase. We next describe each phase in more detail.

#### 3.1.1 Encoder supervised training (Phase I)

The supervised training of the Encoder is illustrated in Fig 3a. Let 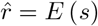 denote the encoded fMRI response from image, *s*, by Encoder *E*. We define an fMRI loss by a convex combination of mean square error and cosine proximity with respect to the ground truth fMRI, *r*. The **fMRI loss** is defined as:

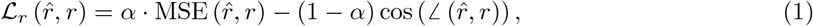

where *α* is a hyperparameter set empirically (*α* = 0.9). We use this loss for training the Encoder *E*.

Notably, in the considered fMRI datasets, the subjects who participated in the experiments were instructed to fixate at the center of the images. Nevertheless, eye movements were not recorded during the scans, thus the fixation performance is not known. To account for the center-fixation uncertainty, we introduced small random shifts (+/− a few pixels) of the input images during Encoder training [13]. This resulted in a substantial improvement in the Encoder performance and subsequently in the image reconstruction quality. Upon completion of Encoder training, we transition to training the Decoder together with the fixed Encoder.

#### 3.1.2 Decoder training (Phase II)

The training loss of our Decoder consists of two main losses illustrated in Fig 3b:

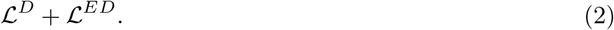

ℒ^*D*^ is a supervised loss on training pairs of image-fMRI. ℒ^*ED*^ (Encoder-Decoder) is an unsupervised loss on unpaired images (without corresponding fMRI responses). Both components of the loss are normalized to have the same order of magnitude (all in the range [0, 1], with equal weights), to guarantee that the total loss is not dominated by any individual component. We found our reconstruction results to be relatively insensitive to the exact balancing between the two-loss components. We next detail each component of the loss.

ℒ^*D*^**: Decoder Supervised Training** is illustrated in Fig 1b. Given {fMRI, Image} training pairs {(*r, s*)}, the supervised loss ℒ^*D*^ is imposed on the decoded image, 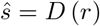, and is defined via the image reconstruction objective, ℒ_*s*_, as

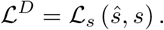

ℒ_*s*_ consists of losses on image RGB values, ℒ_*RGB*_, as well as losses on Deep Image Features extracted from the image using a pretrained VGG16 network [47] (a deep network that was trained for the task of object recognition from images). Using features of pretrained deep neural network classifiers was recently introduced to the task of reconstructing images from fMRI responses in [12, 13], although our implementation is somewhat different (for a comparison of the approaches, see the Discussion section). We denote the deep features extracted from an image, *s*, by *φ* (*s*), on which we apply a ***Perceptual Similarity*** criterion, ℒ_*perceptual*_. This last component gave a significant performance leap. Unlike our preliminary work [45], where we imposed only a Mean-Square-Error loss on the low level features alone (hence failed to capture or exploit any ”semantic” appearance or interpretation), here we impose Perceptual similarity [46] using the outputs of all the five feature-extractor blocks of VGG (from low to high VGG layers, i.e., lower-to-higher “semantic” levels), denoted as 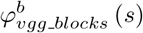 for a particular block *b*. This metric is implemented by cosine similarity between channel-normalized ground-truth and predicted features at each block output. The complete criterion is then a sum of the block-wise contributions. The Image loss for a reconstructed image 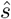 thus reads:

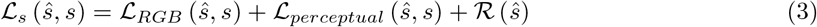

where:

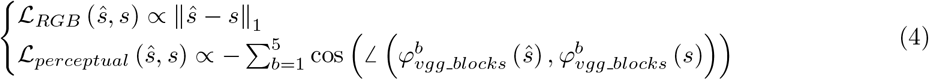

The last term, 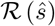, corresponds to total variation (TV) regularization of the reconstructed (decoded) image, 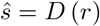. The Image loss in Eq. 3 is also used for the self-supervised Encoder-Decoder training on <u>unpaired images</u> (images without fMRI), as explained next.

ℒ^*ED*^**: Encoder-Decoder training on unpaired Natural Images** is illustrated in Fig 1d. This objective enables to train on any desired unpaired image (images for which no fMRI was ever recorded), well beyond the 1200 or 1750 images included in the fMRI dataset (‘fMRI on ImageNet’ or ‘vim-1’ respectively). In particular, we used ~50K additional natural images from ImageNet’s 1000-class data [44] (excluding the test classes). *This allows adaptation to the statistics of many more novel semantic categories, thus learning the common higher-level feature representation of various novel classes.* To train on images without corresponding fMRI responses, we map images to themselves through our Encoder-Decoder transformation,

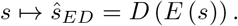

The unsupervised component ℒ^*ED*^ of the loss in Eq. 2 on unpaired images, *s*, reads:

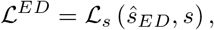

where ℒ_*s*_ is the Image loss defined in Eq 3. In other words, ℒ^*ED*^ imposes <u>cycle-consistency</u> on any natural image, but at a <u>perceptual level</u> (not only at the pixel level). Note that this high-level perceptual consistency does <u>not</u> require using any class labels in the training.

### 3.2 Deep Architecture

We focused on 112×112 RGB or grayscale image reconstruction (depending on the dataset), although our method works well also on other resolutions.

**Architectures** of the Encoder and the Decoder are illustrated in Fig 6. The Encoder comprises four parallel branches of representation, built on top of features extracted from blocks 1-4 of VGG19. This enables to benefit from the hierarchy of “semantic” levels of the pretrained VGG network. The outputs of the four resulting branches (with their various resolutions) are then fed into branch-specific learned convolutional modules, which are designed to reduce the representation’s dimensions to more compact representations of 28×28×32 or 14×14×32 (Height×Width×ConvolutionChannels). These modules consist of batch normalization, 3×3 convolution with 32 channels, ReLU, ×2 subsampling, and batch normalization. The first branch is preceded by an additional ×2 maximum pooling while the fourth branch is not subsampled. Inspired by the feature-weighted receptive field [48] for complex feature spaces, we designed a locally-connected layer which acts on the spatial and channel dimensions separately. This induced separability enables a dramatic decrease in the number of parameters required to regress the voxel activations. In this spatial-feature locally-connected layer, for each spatial coordinate we stack along the channel dimension the values of the immediate 9 neighboring coordinates. This results in tensors of shapes 26×26×288 for branches 1-3 and 12×12×288 for branch 4 (after eliminating the boundaries), and effectively constrains spatial association among adjacent pixels within 3×3 patches. Next, the tensors are multiplied by voxel-specific spatial maps, which learn the voxels’ receptive fields. This outputs space-reduced tensors of shape 288×*N_v_*, where *N_v_* is the number of voxels. To encourage the smoothness of the spatial maps, we penalize for the within-map total variation. The next layer is a cross-channel locally-connected layer, which weighs the contribution of each feature/channel per voxel and outputs a vector of size *N_v_*. This vector is produced by every branch separately. Finally, the outputs of the 4 branches are concatenated along a new dimension and followed by a locally-connected layer (4×*N_v_* parameters) that weighs the contribution from each branch and output the final fMRI activations. We initialized all weights using Glorot normal initializer, except for the last layer, which was 1-initialized (and restricted to remain non-negative).

**Figure 6.**
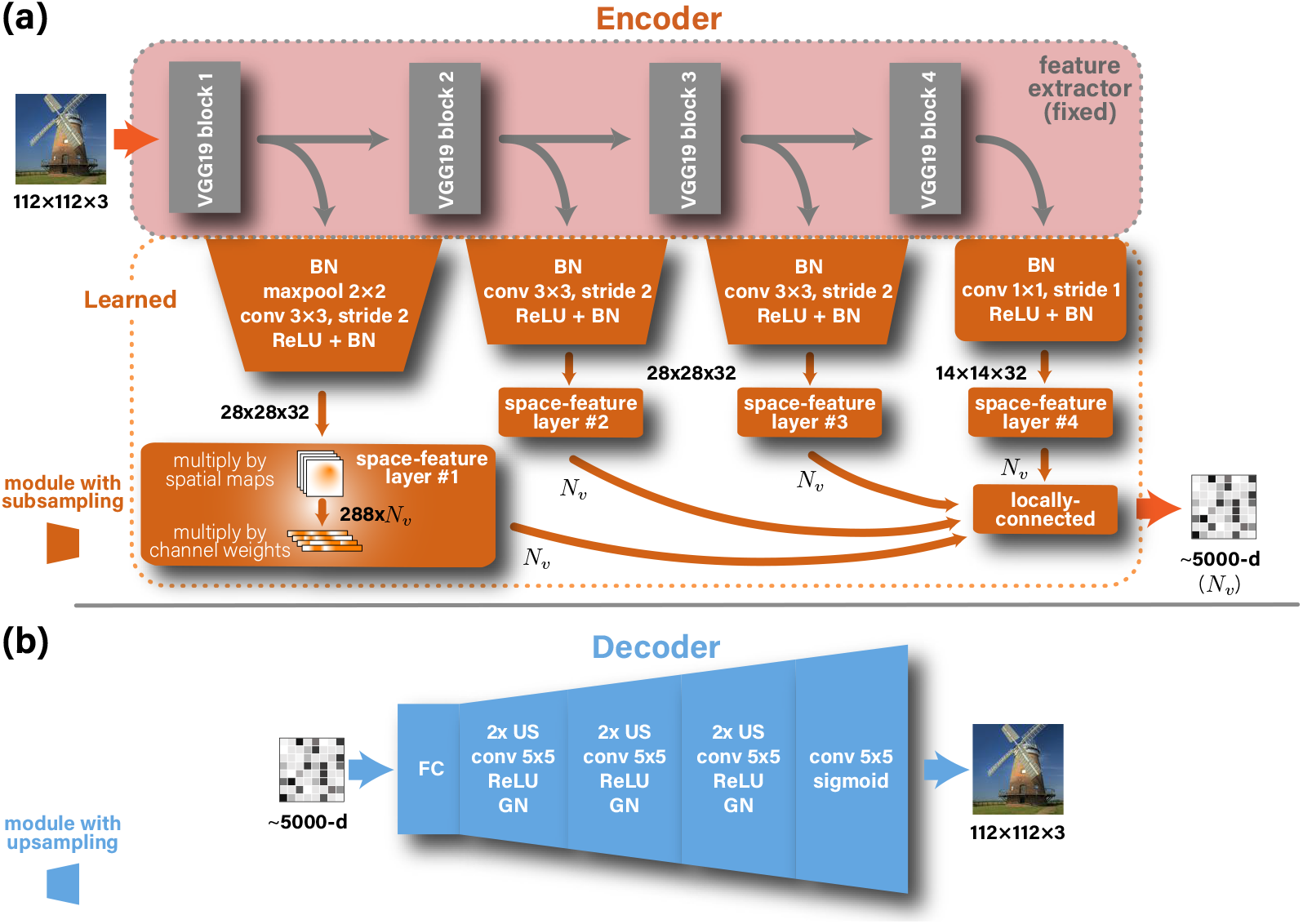
Encoder & Decoder Architectures. BN, GN, US, and ReLU stand for batch normalization, group normalization, up-sampling, and rectified linear unit, respectively. We designed a custom space-feature locally-connected layer (see text).

The Decoder architecture uses a locally-connected layer to transform and reshape the input vector-form fMRI input into 64 feature maps with spatial resolution 14×14. This representation is then followed by three blocks, and each consists of: (i) ×2 up-sampling, (ii) 5×5 convolution with unity stride, 64 channels, and ReLU activation, and (iii) group normalization (16 groups). To yield the output image we finally performed an additional convolution, similar to the preceding ones, but with three channels to represent colors, and a sigmoid activation to keep the output values in the 0-1 range. We used Glorot-normal [49] to initialize the weights.

### 3.3 Experimental datasets

We tested our self-supervised approach on two publicly available (and very different) benchmark fMRI datasets that we summarize in Table 1. These datasets provide elicited fMRI recordings of human subjects paired with their corresponding underlying natural images. In both datasets, subjects were instructed to fixate at the center of the images. The same architectures and hyperparameters were used for both datasets. In ‘fMRI on ImageNet’ dataset [43], which was used also for image classification, 1250 images were drawn from 200 selected ImageNet categories. 150 categories (classes) were used as training data (8 images per category – altogether 1200 training images). The 50 remaining image categories were designated as the novel test categories, represented by 50 test images (1 image from each test category) to be recovered from their fMRI recordings (“test-fMRI”). Importantly, we averaged across same-stimulus repeated fMRI recordings, when available (Table 1). This gave rise to a higher signal-to-noise ratio fMRI samples, and thus to best reconstruction and classification performance (see Supplementary-Material for single-trial reconstruction, identification, and classification results). This further enabled direct comparison with previous works [12, 13, 36].

#### External (unpaired) images database

For unsupervised training on unpaired images (Encoder-Decoder objective, Fig 1d) we used additional 49K natural images from 980 classes of ImageNet (”ILSVRC”) train-data [44]. We verified that the images and categories in our additional unpaired external dataset do not overlap with the test-images and test-categories in the ‘fMRI on ImageNet’ dataset (the inference target). Since the 50 test classes of ‘fMRI on ImageNet’ [43] partially overlap with the 1000 original ILSVRC classes, we particularly discarded the 20 overlapping classes. At test-time, we tested classification against 1030 classes (980+50).

## 4 Results

To test the feasibility of our approach we experimented with two publicly available benchmark fMRI datasets: (i) fMRI on ImageNet [43], and (ii) vim-1 [26].

### 4.1 Image Reconstruction from fMRI

Fig 2 shows our results with the proposed method, including the combined supervised and self-supervised training with perceptual criteria. These results (in red frames – 3rd column) are contrasted with the results obtainable when using supervised training only (e.g., the 1200 paired training examples of the ‘fMRI on ImageNet’ dataset – 2nd column), as well as when not using perceptual criteria (4th column). All the displayed images were reconstructed from the dataset’s test-fMRI cohort (fMRI of new images from never-before-seen semantic categories). The red-framed images show many faithfully-reconstructed shapes, textures, and colors, which depict recognizable scenes and objects. In contrast, using the supervised objective alone led to reconstructions that were considerably less recognizable (2nd column). Similarly, excluding the perceptual criteria as well as the extended Encoder architecture led to reconstructions that were less perceptually understandable (4th column). The reconstructions of the entire test cohort (50 images in the ‘fMRI on ImageNet’ dataset) as well as an assessment of the separated contribution of the perceptual criteria and the extended Encoder architecture can be found in the Supplementary-Material.

To verify that our method can successfully be applied to different subjects, Fig 7 shows the reconstructions for all five subjects in the ‘fMRI on ImageNet’ dataset. Note that using fMRI data of different subjects give rise to varying quality of reconstruction as driven by the varied SNR in the subjects’ fMRI data (which is substantially affected by the subject’s ability to maintain fixation at the center pixel in all repeated scans of the same image). Nevertheless, clear and common identifying markers of the ground truth image appear across all subjects. The red frame indicates the results for the best subject (Subject 3 in the dataset) which is the subject of focus in the remaining parts of this paper (unless mentioned otherwise).

**Figure 7.**
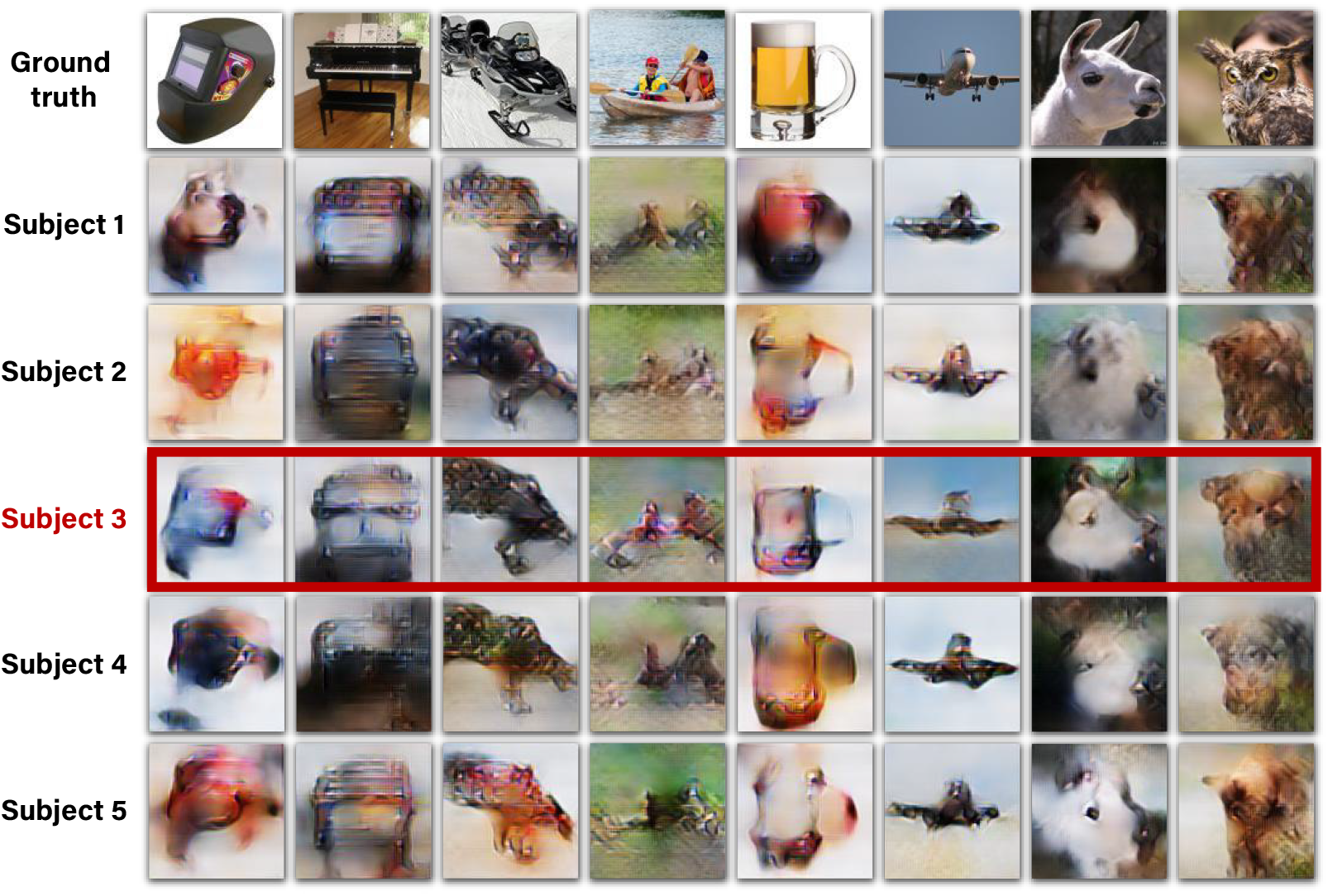
Reconstructions for all five subjects in ‘fMRI on ImageNet’ [43]. Reconstructed images when using the full method, which includes training on unpaired data (1d,e). Reconstruction quality varies across subjects, depending on noise-ceiling/SNR of subjects’ data (voxel median noise-ceiling for subjects 1-5: 0.56, 0.57, 0.73, 0.68, 0.58). Subject 3 (in the dataset), which is framed above in red, is the subject of focus in the remaining parts of this paper unless remarked otherwise.

#### 4.1.1 Comparison with state-of-the-art image reconstruction methods

We compared our reconstruction results against the two leading methods: Shen et al. [12]) and St-Yves et al. [36] – each on its relevant dataset. Fig 8a,b compares the results of our method with those two methods (both of which are deep-learning GAN-based methods). Visual comparison of [12, 36] with our method (Fig 8a,b) highlights that despite their natural-like visual appearance enforced by the GAN, the reconstructed images of [12, 36] are often not faithful to the underlying ground truth image. We further report quantitative comparisons, both by image-metric-based evaluation and by human visual evaluation for the top-SNR subject from each dataset (Subject 3 in ‘fMRI on ImageNet’ dataset [43]; Subject 1 in ‘vim-1‘ dataset [26]). Our quantitative evaluations are based on an n-way identification task [12, 31, 33, 37]. Namely, each reconstructed image is compared against *n* candidate images (the ground truth image, and (*n* − 1) other randomly selected images), and the goal is to identify its ground truth. We considered two identification methods under this task:

**Figure 8.**
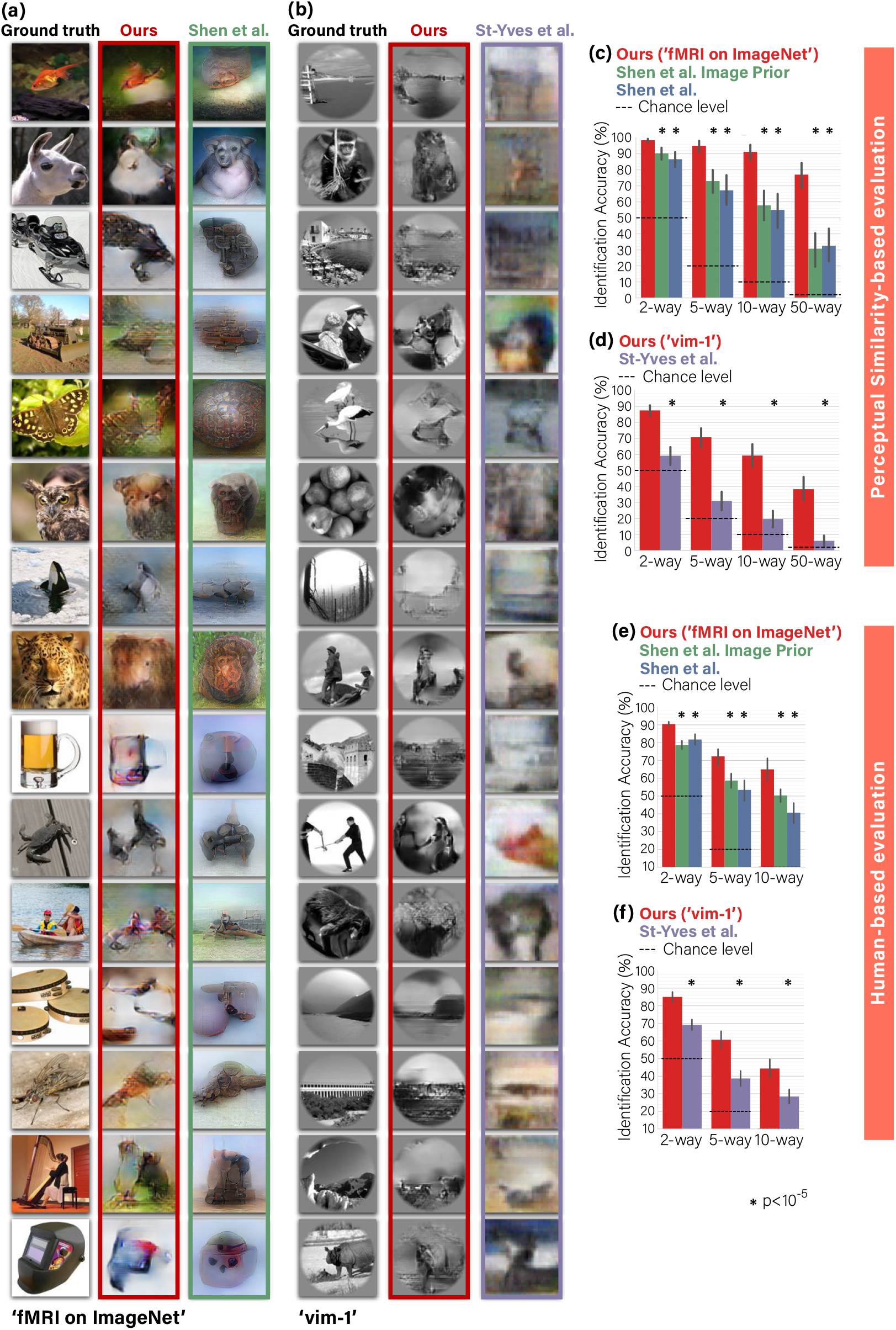
Comparison of image-reconstruction with state-of-the-art methods. **(a), (b)** Visual comparison with [12, 36] – each compared on its relevant dataset. Our method reconstructs shapes, details and global layout in images better than the leading methods. **(c), (d)** Quantitative comparisons of identification accuracy (per method) in n-way identification task according to Perceptual Similarity metric (see text for details). **(e), (f)** n-way identification responses of human raters via Mechanical Turk. Our self-supervised approach significantly outperforms all baseline methods on two datasets and across n-way difficulty levels by both types of experiments – image-metric-based and behavioral human-based (Wilcoxon, N = 50, 120 for panels **(c), (d)**; Mann-Whitney, N = 45 for panels **(e), (f)**). 95% Confidence Intervals by bootstrap shown on charts.

##### (i) Image-metric-based identification performed using the Perceptual Similarity metric [46] (between the reconstructed image and the candidate image)

Each reconstructed image is compared against *n* images, and the nearest-neighbor candidate image under this metric was determined to be the identified ‘correct’ image. Panels 8c,d show the correct-identification rate (for each method separately) for n-way identification tasks for *n* = 2, 5, 10, 50. We evaluate our method and two variants of the method provided by [12] on the ‘fMRI on ImageNet’ benchmark dataset (Fig 8c). In 2-way identification task our method scored an accuracy of 98.5% (*SEM* ^1^ = 0.6%, *N* = 50). Our accuracy remains quite high for all *n*, outperforming both variants of [12] by a margin of 8-46% across all task difficulty levels (*n* = 2, 5, 10, 50). We repeated the analysis for ‘vim-1’ fMRI dataset (Fig 8d), where our method scored an accuracy of 87.5% (*SEM* = 1.7%, *N* = 120) (2-way task), outperforming the method from [36] by a large margin of 28-40% across the same difficulty levels. Particularly in the challenging 50-way task our method achieved striking leaps: outperforming the baseline by more than x2 prediction accuracy in ‘fMRI on ImageNet’, and more than x10 better prediction accuracy in ‘vim-1’. Importantly, the statistical power of these findings generalizes beyond the specific 50 test examples (image-fMRI) in ‘fMRI on ImageNet’, or the 120 test examples in ‘vim-1’ (Wilcoxon test, *N* = 50, 120, *p* < 10^−5^).

##### (ii) Human-based identification

Panels 8e,f show reconstruction evaluation results when repeating the same quantitative comparison approach, but this time outsourcing the n-way identification task (*n* = 2, 5, 10) to random human raters. We used Mechanical Turk to launch surveys to new 45 raters for each evaluated method. Our method scored 90.4% (*SEM* = 0.6%, *N* = 45) and 85.0% (*SEM* = 1.5%, *N* = 45) in a 2-way identification task on ‘fMRI on ImageNet’ and ‘vim-1’ respectively; Scaling the task difficulty up to 10-way, our method scored 65.0% (*SEM* = 3.0%, *N* = 45) and 44.3% (*SEM* = 2.5%, *N* = 45). Overall our method significantly outperformed the previous methods, on both datasets, and across difficulty levels by a margin of at least 8.7% (Mann-Whitney test, *N* = 45, *p* < .001).

Notably, the identification accuracy of each reconstructed image *when using the image-metric for evaluation*, mostly depended on the reconstruction quality of the specific image, and was robust to randomizing the selection of the (non-ground-truth) candidate images. Furthermore, the choice of candidate images *in the human-based evaluation* was fixed across the raters and the methods we compared. Therefore, while the two types of evaluations (image-metric and human-based) consider a seemingly similar n-way identification task, *they are not directly comparable*. Additionally, note that they suggest different statistical generalization insights – generalization beyond the specific set of test examples, when using the image-metric (Wilcoxon test, *N* = 50, 120, *p* < 10^−5^), and generalization beyond the specific pool of human raters, in the human-based evaluations (Mann-Whitney test, *N* = 45, *p* < .001). Overall, our method significantly outperforms state-of-the-art methods by a large margin in both image-metric-based and human-based evaluations.

### 4.2 Decoding rich novel semantic categories of reconstructed images

The benefit of our self-supervised approach extends beyond the task of image reconstruction. It introduces significant gains in the task of semantic classification of the reconstructed images to their novel semantic categories (i.e., categories/classes never seen during training – classes neither represented in the ‘paired’ training set, nor in the ‘unpaired’ external images). For the classification task, we consider the ‘fMRI on ImageNet’ dataset [43], whose train-set contains 1200 images from 150 ImageNet classes. Its test-set contains 50 images from *disjoint* ImageNet classes. The unpaired external images used for the self-supervised training of our network, are drawn from 980 ImageNet classes. Notably, the unpaired images have no fMRI recordings whatsoever. The 50 test classes (to be recovered from the 50 test-fMRIs), are neither included in the 150 ’paired’ train-classes, nor in the 980 classes of the ’unpaired’ external images. *These are totally novel classes* (details in Sec 3.3). When checking the classification results of the 50 test fMRI, we test the classification of their reconstructed images against the gallery of 1030 ImageNet classes^2^. As mentioned earlier (Sec 2.2), such classification requires no explicit class-labels in the training (see Fig 4). It is achieved by comparing the Deep-Features of the reconstructed image, with the Deep-Feature class-representative vector – one representative vector for each of the 1030 classes in our gallery (or any other gallery of novel classes).

Fig 5a exemplifies novel-class classification results by our method. We present each reconstructed image alongside the five top predicted (‘Top-5’) classes among the 1030 classes. For visualization purpose only, each of the Top-5 classes are visually exemplified by the nearest-neighbor image (most similar Deep-Features) among <u>100 randomly selected images</u> from that class label. The first five rows show correct-classification cases at Top-5 accuracy level. In these cases our method successfully includes the ground truth class among the nearest five classes (marked by a red frame). Interestingly, many of the non-ground-truth classes which are assigned among the Top-5, are also reasonable, frequently representing semantic and visual content, which is reminiscent of the ground truth class as well. For example, the ‘housefly’ is found similar also to other flies and comparable shape insects, and the ‘beer mug’ is also found similar to other types of mugs. The bottom rows show incorrect-classification cases, where the ground truth class is not found among the Top-5 classes. Nonetheless, even in these allegedly “failure” cases, many of the Top-5 classes (and even Top-1) are considerably relevant both semantically and visually. For example, the reconstructed ‘video’ image was associated with the ‘television’ class; the ‘duck’ was wrongly classified as an ‘ostrich’ ; the ‘leopard’ was wrongly classified as a ’lion’; the ‘minaret’ was associated with several other tower-like classes, etc.

We evaluate the performance of our classification results using two types of quantitative evaluations: (i) the “Classification Rank” – the average rank of the correct class among all classes (Table 2), and (ii) the more familiar “n-way classification” accuracy (evaluated for *n* = 50; 100; 500; 1000, Fig 5b). These 2 types of numerical evaluation are detailed below.

#### (i) Classification Rank

As explained in Sec 2.2, the location of the ground-truth class within the list of 1030 classes (sorted according to our similarity-based classification measure - see Sec 2.2), provides a “rank score” for evaluating our classification accuracy per image (where rank=1 out of 1030 means perfect score). Moreover, this rank criterion further allows to asses the *generalization power* of our method beyond the limited test-set (the 50 test images and their 50 specific test classes available in the benchmark dataset). Table 2 summarizes novel-class classification rank results for all five subjects in ‘fMRI on ImageNet’. To demonstrate the power of our self-supervised approach, we compare its classification performance with a baseline of the purely supervised approach. This baseline uses reconstructions that were produced by a Decoder trained using the scarce paired data alone (Fig 1b). This comparison shows a leap improvement in median classification rank in favor of our self-supervised approach in all five subjects. Notably, for Subjects 2-5 (excluding Subjects 1 who has the lowest median noise-ceiling), the advantage of our self-supervised approach generalizes beyond the specifically chosen 50 test images and classes of the considered dataset.

#### (ii) n-way Classification

In addition to the Ranking-score within the 1030 gallery classes (for each test-fMRI), we present another alternative way of evaluating the classification results – through classification accuracy in n-way *classification* experiments (for *n* = 50, 100, 500, 1000), using our automated Deep-Features class-similarity criterion. Fig 5b shows Top-1 classification accuracy across a range of classification task difficulties (shown for Subject 3; the remaining subjects can be found in the Supplementary-Material). The tasks differ in the number of candidate classes (n-way) from which prediction is made (i.e., the percent of cases that the Top-1 predicted class out of *n* class labels is indeed the ground-truth class). Our full method scores 29.9% Top-1 accuracy (*SEM* = 0.3%, *N* = 25000) in 50-way classification task. Even when scaling to 1000-way (as in ImageNet classification), our method scores 12.1% Top-1 accuracy (*SEM* = 0.2%, *N* = 25000), which exceeds chance level accuracy by more than 100-fold. Contrasting this performance with the baseline of the supervised approach shows a striking leap improvement in classification-accuracy in favor of our self-supervised approach: between x2 and x3 accuracy improvement, in all the n-way experiments (*n* = 50, 100, 500, 1000).

We further performed an *ablation study* of the Perceptual Similarity for the task of semantic classification. Fig 5c shows 1000-way Top-5 classification accuracy by our self-supervised method, where the reconstructions used are produced by ablated versions of the Perceptual Similarity [46]. Specifically, we limit the Perceptual Similarity criterion, which is used in Decoder training, to a varying range of Deep VGG layers, starting from using only the outputs of the first block (low ”semantic” features) of VGG16, and up to aggregating outputs from all five blocks (high ”semantic” features) of the pretrained network as in the full method. We find that the classification accuracy of the reconstructed images shows an increasing trend with the number of higher-level features, which are used as reconstruction criteria. This highlights the significance of the Perceptual Similarity reconstruction criterion, which includes higher-level features, for semantic classification. Note that the increasing trend appears to a various degree for different subjects, depending on experiment noise and subject-specific noise-ceiling (e.g., Subject 1 having the lowest noise-ceiling); Nevertheless, the trend is well illustrated by their cross-subject mean accuracy. Furthermore, we find that our classification approach, which is applied onto the reconstructed images, outperforms a direct-fMRI classification approach, where the test-fMRI samples are directly classified (see Supplementary-Material).

Our classification approach is inspired by [43]. Both methods use deep-feature embeddings to search for the nearest class centroid in a gallery of novel classes (our embedding is extracted from the reconstructed image (Fig 4b), whereas that of [43] employs an intermediate deep image-embedding decoded from fMRI). Notably, [43] presented the classification of novel categories in a 2-way task (i.e., discriminating between the correct category and a single random category). Here we scale up this classification task to 1000-way (i.e., finding the correct category among 1000 rich categories). *To the best of our knowledge, we are the first to demonstrate such large-scale semantic classification capabilities from fMRI data*.

### 4.3 Predominance of early visual areas in reconstruction

To reconstruct the images of ‘fMRI on ImageNet’, we considered 4600 visual-cortex voxels provided and labeled in [43]. We studied the contribution of different visual areas to our reconstruction performance. To this end, we selected subsets of voxels according to their marked brain areas, and restricted the training of our Encoder/Decoder to those voxels. Fig 9 shows reconstruction results when using voxels only from the following visual areas: (i) V1 (870 voxels), (ii) V1-V3, which refer to as Lower Visual Cortex (LVC, 2300 voxels), (iii) Fusiform Face Area (FFA), Parahippocampal Place Area (PPA), and Lateral Occipital Cortex (LOC), which we refer to as Higher Visual Cortex (HVC, 2150 voxels), or (iv) Full Visual Cortex (VC = LVC + V4 + HVC, 4600 voxels). These results show that the early visual areas, particularly V1-V3 (LVC), contain most of the information recoverable by our method. Considering voxels only from HVC leads to substantial degradation in performance despite comprising approximately half of the complete visual cortex voxels. Nevertheless, the higher visual areas clearly add semantic interpretability to the reconstructed images (which is evident when comparing the reconstructions from the Full VC, to those from LVC only).

**Figure 9.**
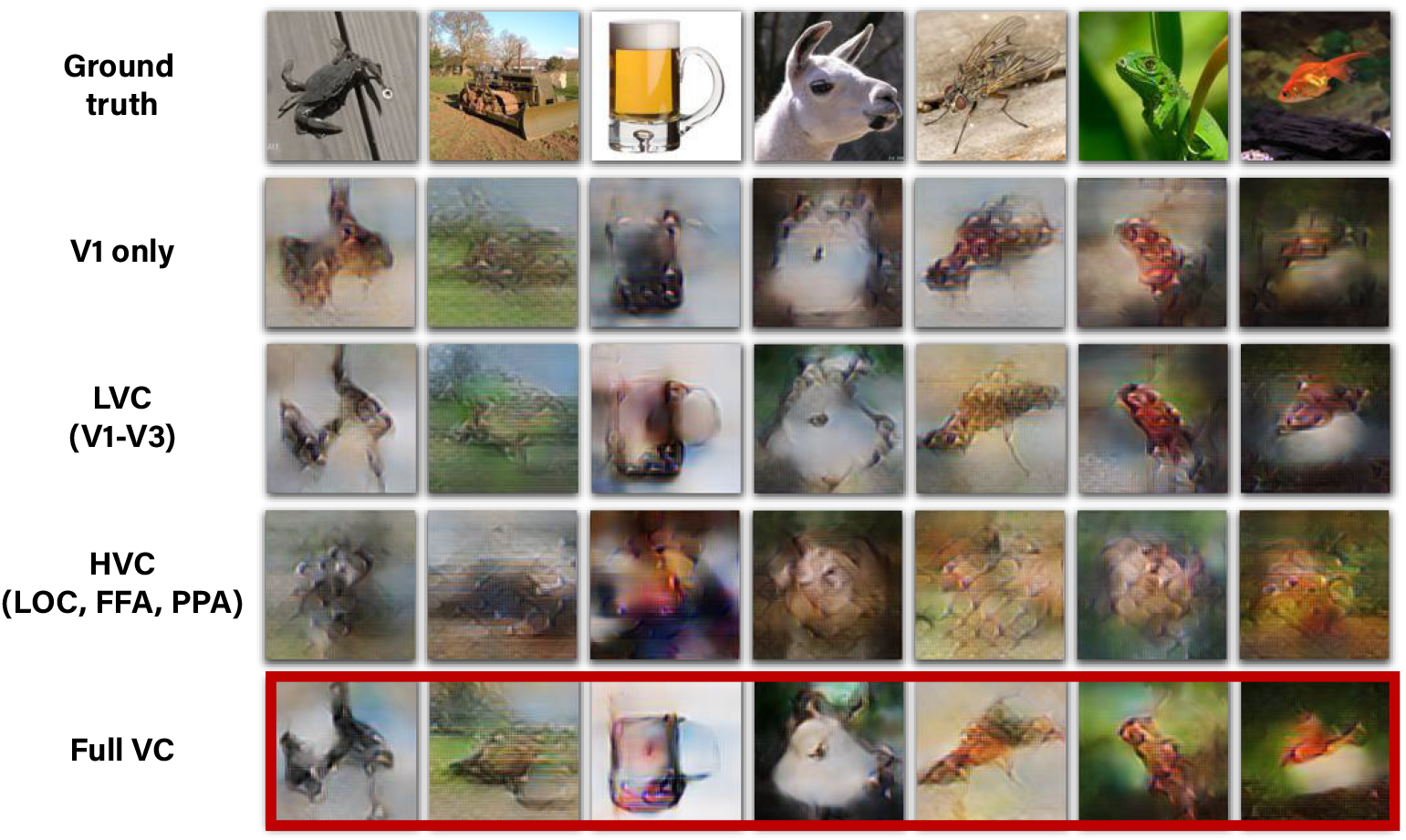
Decoding quality is dominated by early visual areas. Columns show reconstructions using our method with fMRI data from various ROIs in the visual cortex including: • **Primary Visual Cortex** – V1 • **Lower Visual Cortex** – V1-V3 • **Higher Visual Cortex** – Fusiform Face Area (FFA), Parahippocampal Place Area (PPA), Lateral Occipital Cortex (LOC) • **Full Visual Cortex** – LVC + V4 + HVC (in red frame).

Importantly, we found that removing any single visual area from our dataset, including V1, does not degrade the results significantly, suggesting the existence of information redundancy across visual areas. The results are strongly affected only when several regions, specifically the entire early visual cortex, are discarded. Furthermore adding V4 to either LVC or HVC did not change the results significantly.

We studied two extensions for the ROI-specific reconstruction results: (i) We controlled for the ROI-voxel count bias by sampling an equal number of voxels from each ROI. This control did not introduce major changes in the cross-ROI comparison or to the conclusion of early-visual-areas predominance; (ii) We analyzed the ROI-specific contribution also in semantic classification. Our results were consistent with those from the ROI-specific reconstruction experiments, showing predominance of early visual areas also in semantic classification. These results can be found in the Supplementary-Material.

### 4.4 Emergent biologically consistent population receptive fields

The human visual system is characterized by the well-known primate *retinotopic organization* [50–58]. Retinotopy maps reflect the spatial tuning of cortical hypercolumns or of their aggregation into Population Receptive Field (pRF) as in the case of voxel data. Here, we sought to visualize receptive field of voxels as captured by our Encoder & Decoder. To this end we considered two approaches: (i) Interpreting model weights that associate each voxel with spatial locations as a heatmap, and (ii) A gradient-based approach on the Encoder’s input space. While this approach is applicable to the Encoder only, it is favorable over the former approach because it provides a higher resolution receptive field map (i.e., 112×112, following Encoder’s *input dimensions* vs. 26×26 or 14×14, following the layer’s architecture in the Encoder/Decoder, see details in Sec 6). Since both methods gave rise to well aligned receptive fields (per voxel, see Supplementary-Material), we henceforth focused our analysis on the receptive fields recovered for the trained Encoder using the gradient-based approach.

Fig 10a shows receptive fields for several selected voxels, which indicates their <u>spatial locality</u> within the image. Next, we estimated the pRFs eccentricity and polar angle for each voxel. Lastly, we plot these data on the subject-specific cortical surface. Fig 10b,c shows the resulting tuning maps revealing the expected retinotopic organization. This includes the emergence of horizontal and vertical meridians and their transitions, contra-laterality and up-down inversion, and fovea-periphery gradual transition. We emphasize that this organization automatically emerged on its own from our training method involving natural images and fMRI. No biological atlas or other retinotopic prior was imposed. These results support the biological consistency of our model’s predictions.

**Figure 10.**
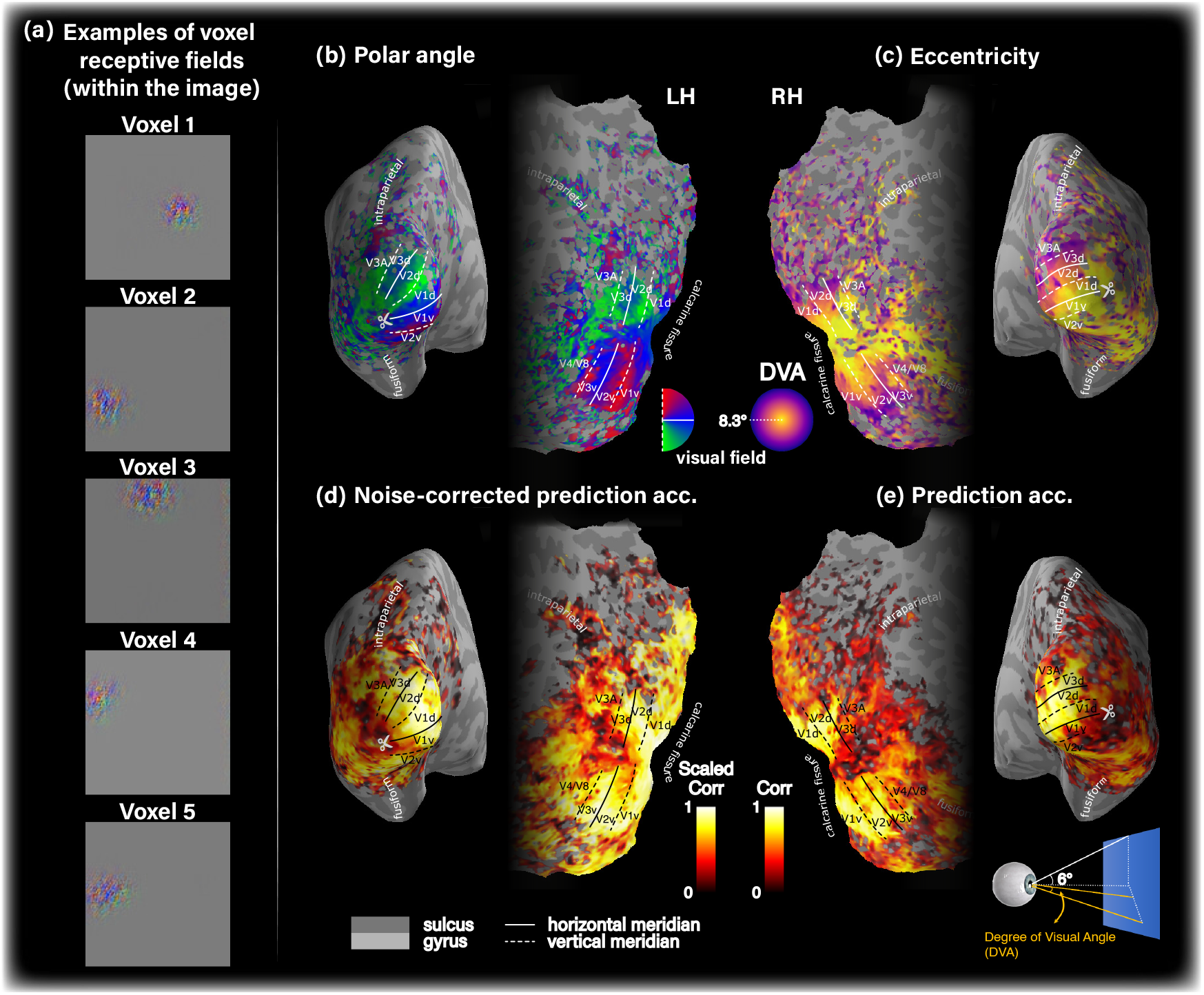
Our models capture biologically consistent voxel tuning properties. **(a)** Receptive field of five selected voxels with high SNR from early visual cortex, which indicates their spatial locality in the image. Panels **(b)-(e)** show single subject data on the corresponding subject-specific cortical surface. **(b)** Polar angle. **(c)** Eccentricity tuning, measured by degree of visual angle (DVA). **(d)** Noise-corrected prediction accuracy. **(e)** Prediction accuracy (non-scaled Pearson correlation). For simplicity we show the data on either left or right hemisphere. Voxel noise-ceiling is coded by transparency level (alpha channel) in all cortical maps.

We sought to analyze the prediction accuracy achieved by of our models. Fig 10d,e show the prediction accuracy distribution (Pearson correlation) of the modeled voxels when normalized by voxel noise-ceiling (Fig 10d) and when not normalized (Fig 10e). The prediction noise-ceiling is used to provide an estimate of the best possible prediction accuracy obtainable given infinite data [23]. These panels show high prediction accuracy in LVC, and low in HVC. Furthermore, they show that throughout the visual cortex, our model markedly saturates the noise-ceiling of the given data. This indicates the sufficient expressive power of our multi-layer encoding model, enabling it to capture the given data complexity. A comparison of the model’s prediction accuracy with a single-layer Encoder architecture from [45] can be found in the Supplementary-Material.

## 5 Discussion

We introduced a novel approach for self-supervised training of image decoders on external, image-only datasets. We show that our method gives rise to both state-of-the-art results in image-reconstruction as well as large scale semantic categorization from fMRI data. To date, the performance in the task of natural image reconstruction and semantic categorization from human fMRI recordings is limited by the characteristics of fMRI datasets. In the typical case, the paired training data are scarce, and represent a narrow semantic coverage.

Our self-supervised training on tens of thousands of additional unpaired images from wide coverage, combined with high-level Perceptual Similarity constrains, adapts the decoding model to the statistics of natural images and novel categories. Our framework enables substantial improvement in both image reconstruction quality <u>and</u> classification capabilities compared to methods that rely only on the scarce paired training data. This self-supervised perceptual approach leads to state-of-the-art image reconstructions from fMRI, of unprecedented quality, as supported by image-metric-based evaluations, as well as extensive human-based evaluations. We accomplish this for two substantially different fMRI datasets using a single method (with the same hyperparameters).

Our self-supervised training on tens of thousands of unpaired external images further leads to unprecedented capabilities in the semantic classification of fMRI data (and moreover, of classes never encountered during training). We consider the challenging 1000-way semantic classification task, and demonstrate a striking leap improvement (more than 2x) in classification performance when applying our self-supervised approach over a purely supervised approach. To the best of our knowledge, we are the first to demonstrate such large-scale semantic classification capabilities (1000-way) from fMRI data.

The improved reconstruction quality achieved when minimizing deep image representation error rather than low-level pixel-based one (Fig 2) supports reports in earlier works that employed similar approaches [12, 13]. In [12], Perceptual Similarity was used to drive image reconstruction by iterative optimization at *test time*. Here, however, the Decoder itself is trained to minimize the higher-level reconstruction error, allowing a non-iterative feed-forward reconstruction at test-time. In this respect, our approach is more compatible with [13], albeit our reconstruction loss is based on the complete Perceptual Similarity metric [46], defined via features extracted *at multiple intermediate layers* of a pretrained object classification network rather than only its almost last layer. Furthermore, this work goes beyond [12, 13] in two aspects: (i) Our Decoder benefited from applying Perceptual Similarity to substantially more data with no fMRI recording. The combined effect of this type of reconstruction criterion with the additional unpaired data is very effective, as evident in Fig 2; (ii) We show that applying Perceptual Similarity not only results in an improved reconstruction quality, but also gives rise to a higher classification accuracy of novel ImageNet classes (classes that were not represented in any of the training data). An important question for future research is whether a Natural Image Prior, such as the Deep Generator Network employed by [12] can be combined with the self-supervised approach in a way that benefits from both the self-supervision on additional unpaired data and the imposed narrower Natural Image search space.

Our ablation studies indicate that reconstruction quality is dominated by data originating from Lower Visual Cortex (V1-V3). The extended architecture of the Encoder, which incorporates high-level deep features, was designed to improve information-harnessing from the Higher Visual Cortex (HVC) as well. Indeed prediction accuracy maps show that the noise-ceiling is saturated throughout the visual cortex, including in higher visual areas. These findings suggest a reasonable representation of the HVC by our model. Nevertheless, the SNR of the data arising from these areas renders them weaker contributors to overall reconstruction quality.

We provide evidence for the retinotopic organization implicitly learned (on its own) by our image-to-fMRI Encoder. This suggests that our models are biologically meaningful, as opposed to tailored and overfit to a limited dataset. Note that while we show data for the Encoder, we verified in our experiments that model voxels in the Decoder and the Encoder indeed agree (while not explicitly forced to do so).

The proposed method currently focuses on data from individual subjects. A natural extension of the present work is to combine information across multiple subjects (see [59]). This is part of our future work.

## 6 Additional Technical Details

### Hyperparameters

We trained the Encoder using Adam optimizer for 50 epochs with an initial learning rate of 1e-3, with a 90% learning rate drop using milestones (20, 30, and 35 epochs). During Decoder training with supervised and unsupervised objectives, each training batch contained 16 pairs (supervised training), and 16 unpaired natural images (randomly sampled from the external image database – images without fMRI). We trained the Decoder for 150 epochs using Adam optimizer with an initial learning rate of 1e-3, and 80% learning rate drop after every 30 epochs.

### Runtime

Our system completes the two-stage training within approximately 1.5 hours using a single Tesla V100 GPU. Once trained, the inference itself (decoding of a new fMRI) is performed in a few milliseconds per image.

### Behavioral experiments

The participants in the Mechanical Turk behavioral experiments gave their online informed consent to be recorded, and were granted financial incentives for every completed survey. The research protocol was reviewed and approved by the Bioethics and Embryonic Stem Cell Research Oversight (ESCRO) Committee at the Weizmann Institute of Science. In order to assure the validity of the behavioral data (e.g. bot observers, fatigue along the survey), we screened subjects according to their score in interleaved sanity check experiments. The sanity check experiments comprised adding to the actual experiments also 10% unexpected trivial identification tasks of mildly degraded versions of the ground truth images, instead of the reconstructed images. We further discarded subjects with MTurk success-score (reputation) lower than 97%. Each survey consisted of 50 or 20 trials corresponding to the number of test-images comparison in ‘fMRI on ImageNet’ [43] or ‘vim-1’ [26]^3^, all of which were reconstructed using a single particular method. In each trial subjects were presented with a reconstructed image and *n* candidate images, the ground-truth image and *n* − 1 distractor images, and were prompted ”Which image at the bottom row is most similar to the image at the top row?”. To assure task difficulty agreement across subjects and reconstruction methods the set of distractor images was randomly selected for each test-image, but remained fixed across surveys; Our results were insensitive to their re-selection.

### Semantic category decoding

We defined the feature vector underlying the class representatives to be the outputs of block 4 in AlexNet, and used Pearson correlation as the distance metric for ranking class representatives. Our experiments showed that using this intermediate representation level as the embedding of choice yields optimal results for classification.

### Voxel receptive field visualization and estimation

We analyzed the receptive field of our encoding model in terms of simulated retinotopy as previously proposed in [39, 60]. To generate analogous spatial tuning maps for our *modeled* voxels, we estimated the voxel spatial tuning captured by our models. We start by visualizing each voxel’s receptive field (pRF) using the trained Encoder. Our primary approach to this end was gradient-based [61, 62]: Given a random input image, we compute the gradient of a particular voxel with respect to this input. This allows to visualize the image which would drive the maximum change in activity at the target voxel. To produce a heat-map, the values within the resulting gradient-image are squared, averaged across the color-channels, and normalized. Next, we define the pRF center as the center of mass of the *preprocessed* map. The preprocessing was designed to minimize noise effects. It included map smoothing with a Gaussian kernel, *σ* = 3, followed by raising the map values to the power of 10 (emphasizing the areas with higher values). About 15% of the voxels had pRF maps which were not confined spatially around a center of mass, and were thus discarded in subsequent analysis. The remaining 85% voxels were considered in the retinotopy maps. In a secondary approach, we recovered voxel receptive field maps for both the Encoder and the Decoder from the learned voxel weights. For the Encoder, we directly visualized the spatial map weights of the space-feature locally-connected layer. For the Decoder, we reshaped the input fully-connected layer to square spatial dimensions and 64 channels and computed the mean square weights across channels to yield a heat map.

### Cortical surface visualization

We reconstructed and flattened the individual cortical surfaces using FreeSurfer, and plotted the data using a custom extension of PySurfer.

### Noise-Ceiling

We estimated the fMRI prediction Noise-Ceiling by half-split over the test data repeats following [23].

### Statistics

We used Wilcoxon signed-rank (paired) test (two-tailed) for significance testing in the image-metric-based multi-image identification experiments. For the rank-classification experiments, and the (unpaired) behavioral experiments we used Mann-Whitney rank test.

## CRediT authorship contribution statement

**Guy Gaziv:** Writing - Original Draft, Conceptualization, Methodology, Investigation, Formal analysis, Visualization. **Roman Beliy:** Conceptualization, Methodology, Investigation, Software. **Niv Granot:** Conceptualization, Methodology, Investigation, Software (added the Perceptual Similarity to the framework). **Assaf Hoogi:** Resources, Investigation (human-based experiments). **Francesca Strappini:** Resources, Visualization (fMRI preprocessing and brain visualization). **Tal Golan:** Resources, Visualization, Formal analysis (fMRI preprocessing, brain visualization, advised on statistical testing). **Michal Irani:** Conceptualization, Methodology, Investigation, Supervision, Funding acquisition. **All authors:** Writing - Review & Editing.

## Data and code availability statements

We used publicly available fMRI datasets as well as image datasets as follows: fMRI on ImageNet, vim-1, and ImageNet. We will make our code publicly available upon publication.

## Acknowledgments

This project has received funding from the European Research Council (**ERC**) under the European Union’s Horizon 2020 research and innovation programme (grant agreement No 788535).

## Footnotes

1 Standard Error of the Mean.

2 ImageNet consists of <u>15K</u> semantic classes, from which only 1000 classes participate in the ImageNet classification challenge (ILSVRC). In ‘fMRI on ImageNet’ [43] which we use, only 20 out of the 50 test classes are included among the ILSVRC classes. The remaining 30 classes are taken from the larger collection of 15K ImageNet classes. Since our gallery is based on the 1000 ILSVRC classes, at train-time we omit the test classes, resulting in 980 train-classes (= 1000 20). At classification test-time, we add the 50 test-labels to the gallery, resulting in 1030 class labels (= 980 + 50).

3 ‘vim-1’ originally contains 120 test-images, however in the behavioral evaluation we considered only the subset of 20 images that were defined in [36] as test-images

