## Supplementary Material for "Self-Supervised Natural Image Reconstruction and Large-Scale Semantic Classification from Brain Activity"

### Reconstruction results of the complete test cohort

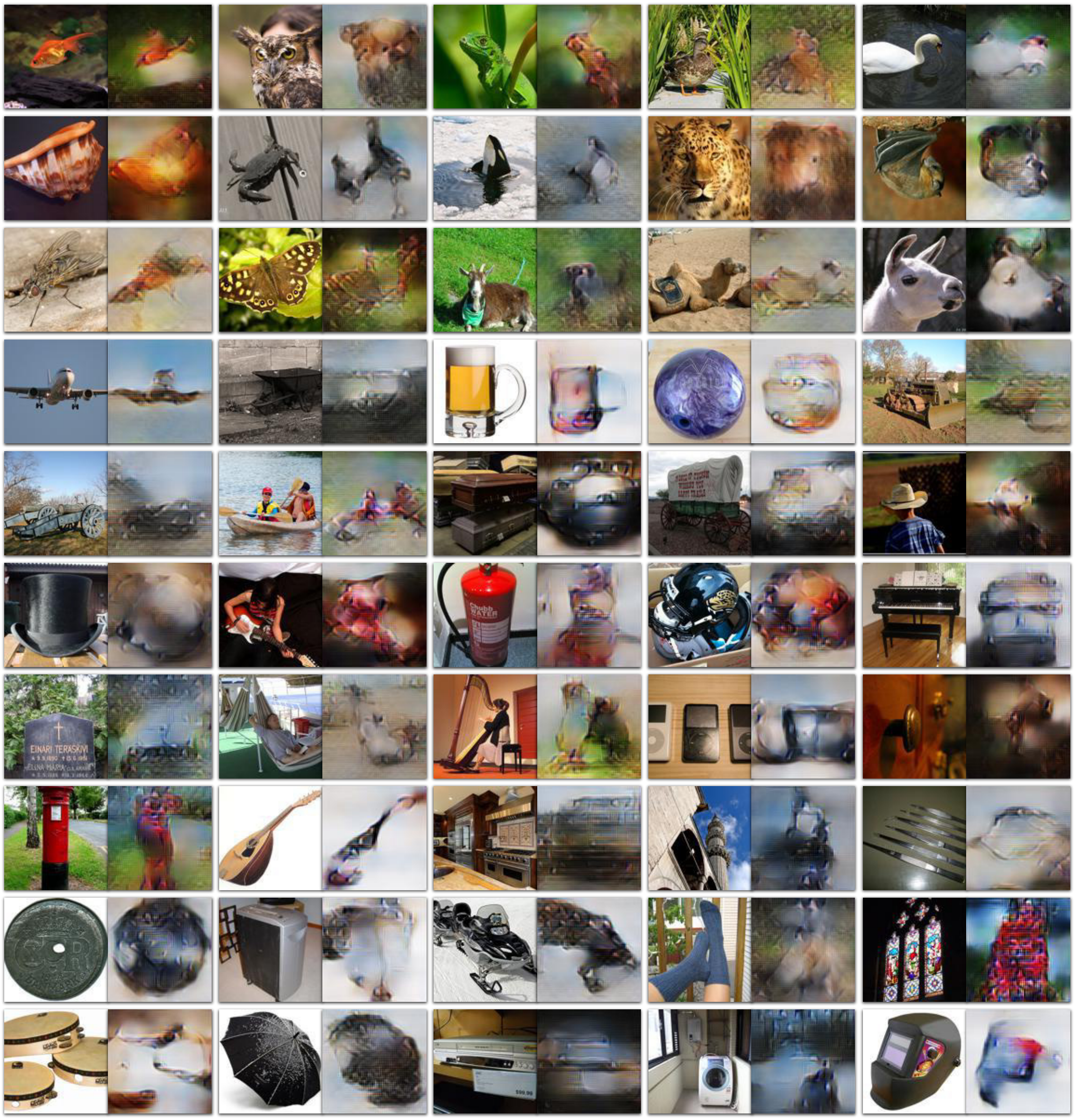


**Figure 1. Collage of reconstructions for the entire 'fMRI on ImageNet' test data (50 images).** Each pair of images shows the reconstructed image (right) side-by-side with its ground truth image (left). We present reconstructions for the subject with the highest noise ceiling, which is Subject 3 in the dataset.

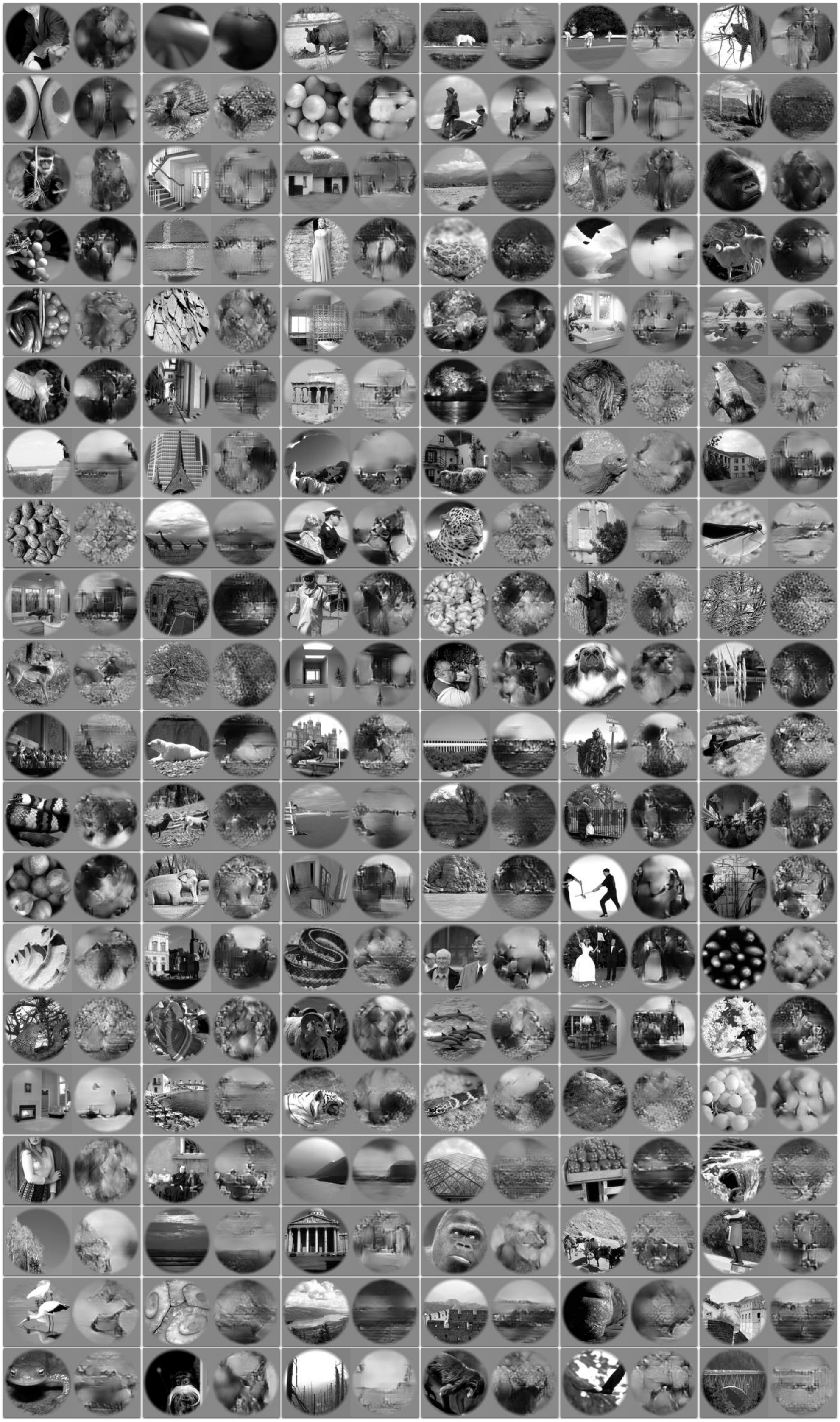

**Figure 2. Collage of reconstructions for the entire 'vim-1' test data (120 images).** Each pair of images shows the reconstructed image (right) side-by-side with its ground truth image (left). The presented reconstructions are for the subject with the highest noise ceiling, subject 1 in the dataset.

### Semantic classification results for other subjects

In the main text we provided Top-1 classification results for subject 3. Fig 3 provides the results obtained for the other three subjects in 'fMRI on ImageNet' [1] with the highest noise-ceiling.

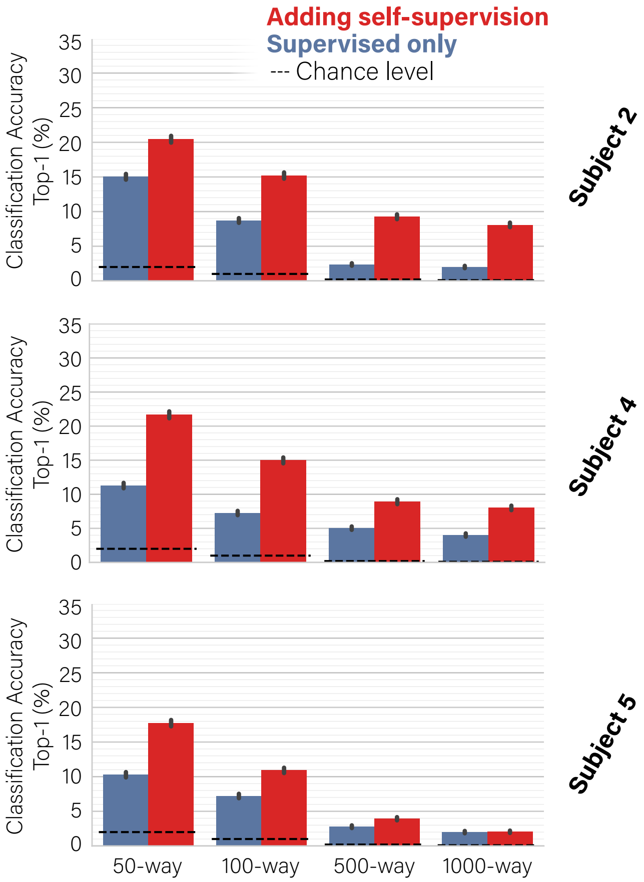


**Figure 3. Self-supervision allows classification to rich and novel semantic categories.** *Mean classification accuracy in an n-way classification task for subjects 2, 4 and 5, up to $n = 1000$, at Top-1 accuracy level. Adding unsupervised training on unpaired data dramatically outperforms the baseline of the supervised approach.*

### Predominance of early visual areas in semantic classification

We extended the ROI-specific analysis also for semantic classification. Fig 4 provides semantic classification results by the complete self-supervised approach. Namely, we computed the classification accuracy using ROI-specific reconstructed images, which were obtained when restricting the training of our Encoder/Decoder to selected subsets of voxels according to their marked visual areas (see Main Text). We present results for the four main subjects in 'fMRI on ImageNet' [1] with the highest noise-ceiling. These results show that the early visual areas, particularly V1-V3 (LVC), dominate our classification performance. Considering voxels only from HVC leads to substantial degradation in performance despite comprising approximately half of the complete visual cortex voxels. Nevertheless, adding the higher visual areas does give rise to significant gains in the semantic classification accuracy. These findings are consistent with those from the ROI-specific reconstruction results, and are reasonable given that our classification method relies on the ROI-specific reconstructed images.

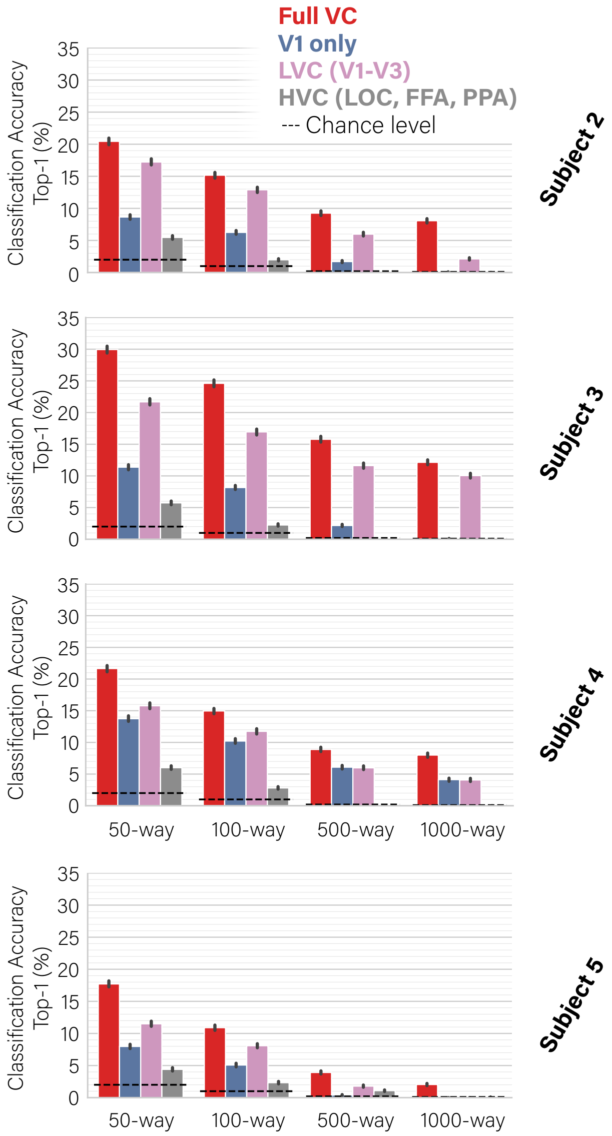


**Figure 4. Predominance of early visual areas in semantic classification.** *Mean classification accuracy in an n-way classification task for subjects 2, 3, 4 and 5, up to* $n = 1000$*, at Top-1 accuracy level. Classification accuracy computed using ROI-specific reconstructed images (see Main Text).*

### Semantic classification directly from fMRI

Our semantic classification approach is based on classifying images that were reconstructed from test-fMRI. We assessed the relative benefit of our approach with a direct-fMRI classification approach, where the test-fMRI samples are directly classified against class representatives. Note that to enable the direct-fMRI approach we had to compute anew the class representatives – in fMRI space – which we performed using our pretrained Encoder.

We find that our reconstructed-image-based classification approach outperforms the direct classification approach. The relative gain is particularly and strongly evident in the challenging large-scale 1000-way classification case. We attribute this gain to two key elements of our approach: (i) It benefits from training on additional unpaired images (the self-supervised approach), and (ii) It benefits from classifying in the high-dimensional and semantically-relevant space defined by a pretrained classification network.

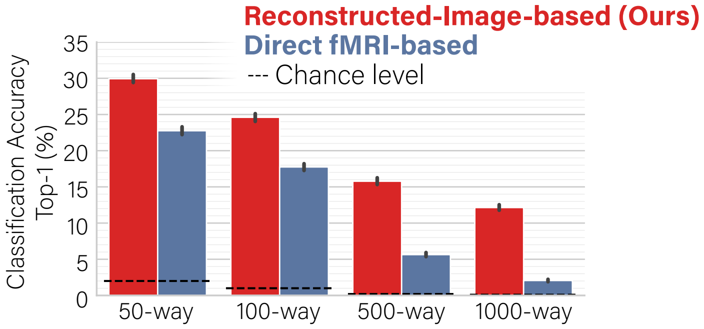


**Figure 5. Reconstructed-Image-based semantic classification outperforms direct fMRI approach.** *Semantic classification results obtained when directly classifying test-fMRI compared with the reconstructed-image-based ones. To enable the former, we re-computed class representatives in fMRI-space using the trained Encoder. Results are demonstrated for 'fMRI onImageNet' [1] Subject 3 (with the highest noise ceiling).*

### Predominance of early visual areas in reconstruction (voxel-count control)

The ROI-specific reconstructions were produced using the varied voxel number of the different ROIs. To account for this potential bias we repeated the analysis when randomly sampling an equal number of voxel from each ROI (including Full VC). We specifically sampled 872 voxels following the minimal voxel count across ROIs (V1). We present these results in Fig 6. We find the reconstructions – and mainly the comparison across the ROI-specific reconstructed images – to be fairly robust to this type of control; We further notice a degradation in the sampled full visual cortex reconstructions.

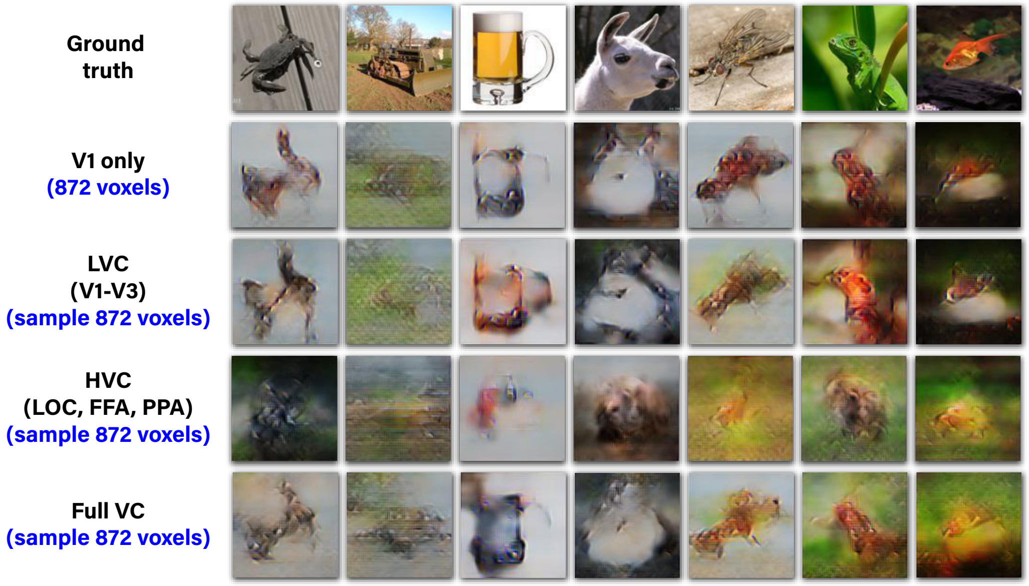


**Figure 6. Decoding quality is dominated by early visual areas (voxel-count control).** *In each ROI we randomly sampled 872 voxels which we then used in our complete self-supervised method.*

### Ablation study: Contribution of perceptual loss and multi-layer Encoder

Fig 7 shows an ablation study to assess the cumulative contributions of the perceptual loss (for training the decoder) and the new multi-layer Encoder. Fig 7a visually demonstrates the dominance of adding the perceptual loss in the improved visual quality over [2]. While replacing the old encoder of [2] with the new multi-layer encoder gives rise to only a moderate extra improvement in the visual quality, the new encoder provides a significant boost to the classification performance, as shown in Fig 7b.

Overall, our findings support the strong significance of adding the perceptual loss for both reconstruction and classification, and suggest the significance of the multi-layer encoder mainly for classification and moderately for reconstruction quality.

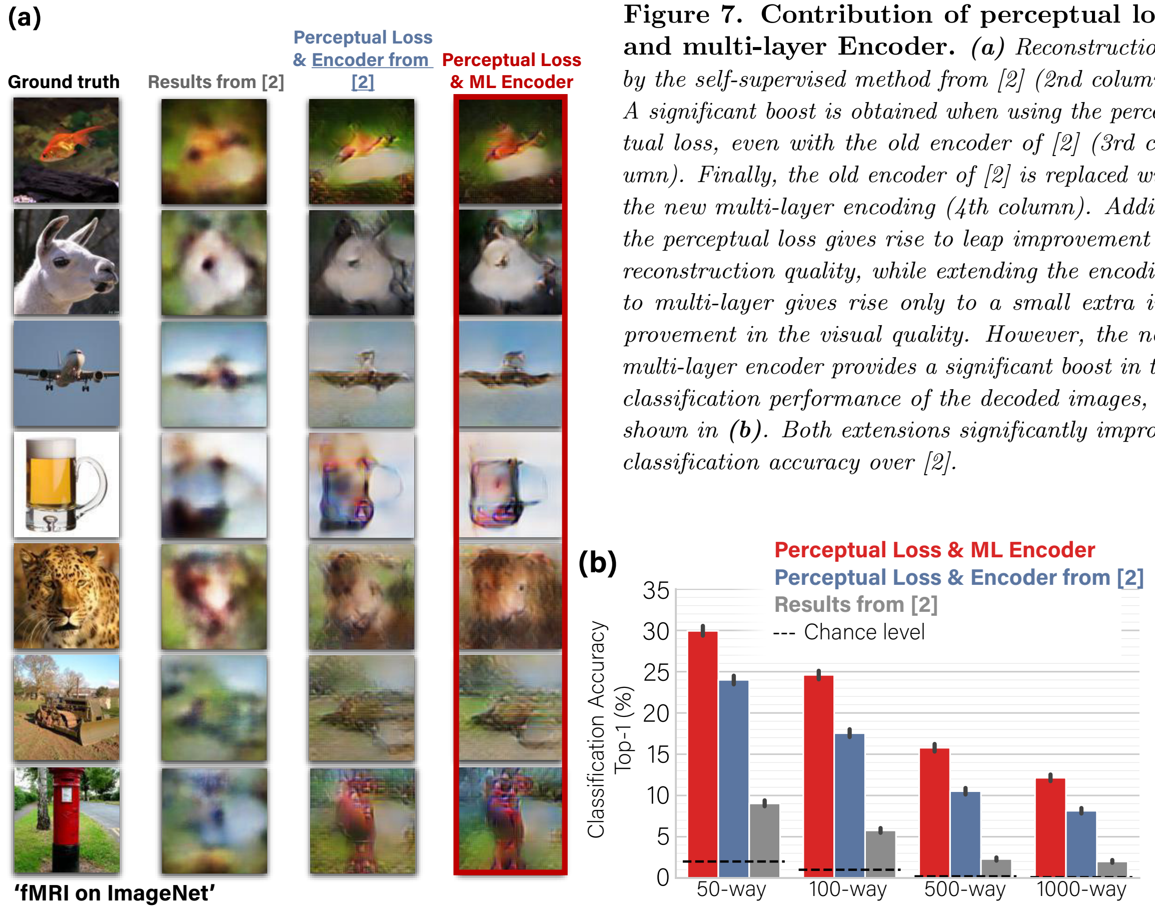


**Figure 7. Contribution of perceptual loss and multi-layer Encoder.** ***(a)*** *Reconstructions by the self-supervised method from [2] (2nd column). A significant boost is obtained when using the perceptual loss, even with the old encoder of [2] (3rd column). Finally, the old encoder of [2] is replaced with the new multi-layer encoding (4th column). Adding the perceptual loss gives rise to leap improvement in reconstruction quality, while extending the encoding to multi-layer gives rise only to a small extra improvement in the visual quality. However, the new multi-layer encoder provides a significant boost in the classification performance of the decoded images, as shown in* ***(b)****. Both extensions significantly improve classification accuracy over [2].*

### Ablation study: Contribution of multi-layer Encoder architecture to fMRI prediction accuracy

Fig 8 shows the per voxel prediction accuracy of our extended multi-layer encoding network compared with the single-layer architecture from [2]. We find that the multi-layer encoding network, which benefits from higher-level semantic features of its pretrained backbone network, is better at predicting voxel activation at higher visual areas.

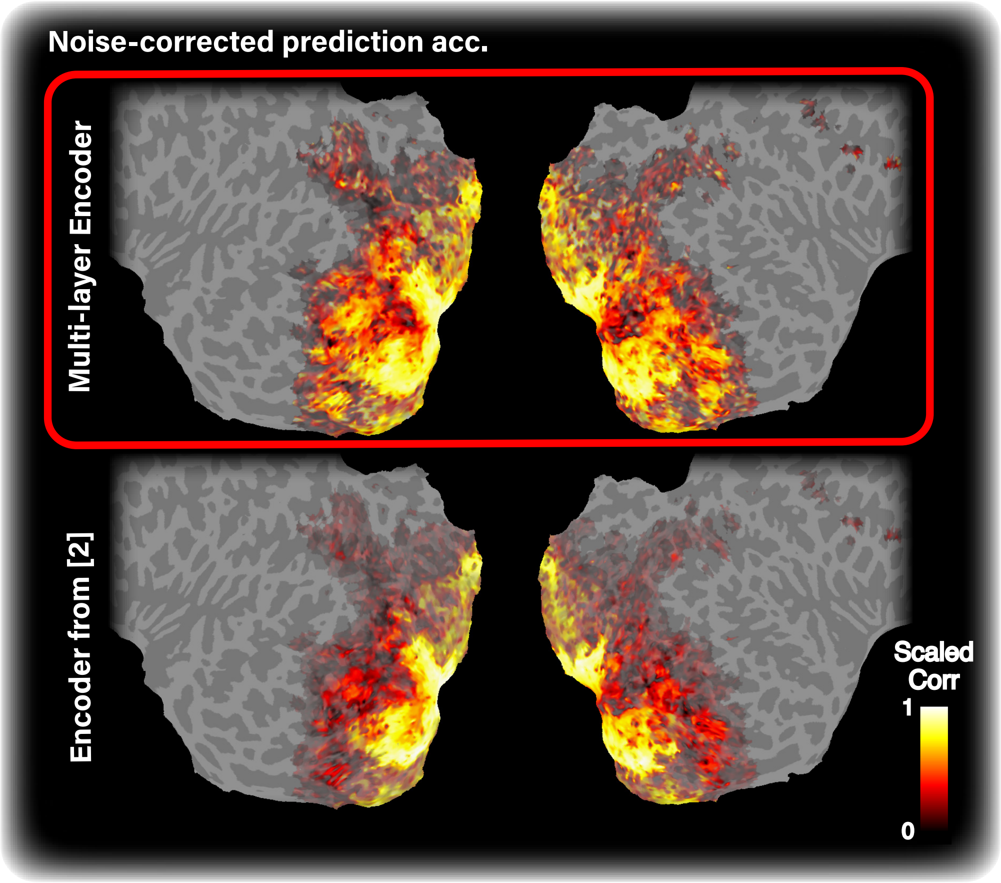


**Figure 8. Contribution of multi-layer Encoder architecture to prediction accuracy.** *Noise-correct prediction accuracy for 'fMRI on ImageNet' [1] Subject 3 (with the highest noise ceiling). Voxel noise-ceiling is coded by transparency level (alpha channel) in all cortical maps.*

### Consistency of voxel receptive field recovery across Encoder/Decoder

We considered two approaches for analyzing voxel receptive fields: (i) Directly interpreting weights that associate a voxel with a spatial location. This approach is applicable to both the Encoder and the Decoder, and provides a spatially coarse resolution receptive field map (i.e., 26x26 or 14x14, following the layer's architecture in the Encoder/Decoder); (ii) A gradient-based approach, which is applicable to the Encoder only, but provides a high resolution receptive field map (i.e., 112x112, following Encoder's *input dimensions*).

Fig 9 shows heat maps of the weight-based receptive fields (Approach (i)) for a subset of voxels for both the Encoder and the Decoder. Furthermore, we present the high resolution gradient-based receptive field maps (approach (ii)) for the Encoder given random noise input. These results indicate the consistency across the two methods as well as the alignment across the encoding and the decoding models.

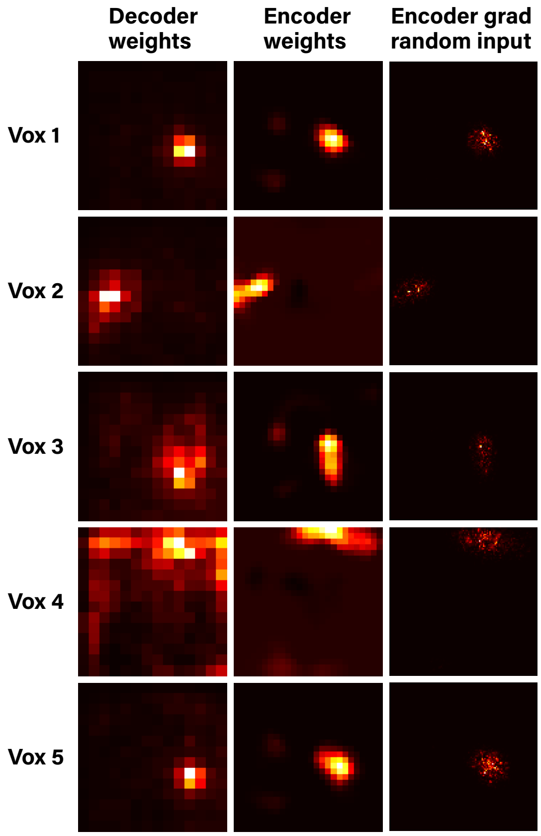


**Figure 9. Voxel receptive field is consistent across Encoder/Decoder and across visualization methods.** *Visualization of receptive field for five selected voxels (with high SNR). The two leftmost columns show receptive field in the Decoder and the Encoder, as reflected by their learned layer weights. The rightmost column shows receptive field in the Encoder as reflected by voxel-activation gradient w.r.t. image pixel values, where the initial input was a random image.*

### Single-trial reconstruction, identification, and classification

We used averaged fMRI data across repeated trials in all of the analyses described in the main text. Specifically, in the fMRI on ImageNet [1] version used, we averaged across the 35 test-fMRI repeats (no repeats exist for train-fMRI). Fig 10 provides examples of reconstructed images from single-trial fMRI data and shows identification and classification results using single-trial reconstructed images. We find that using the single-trial test data significantly degrades performance across the board. Specifically, it gives rise to high-variance reconstructions across repeats and nearly abolishes the large-scale classification capability.

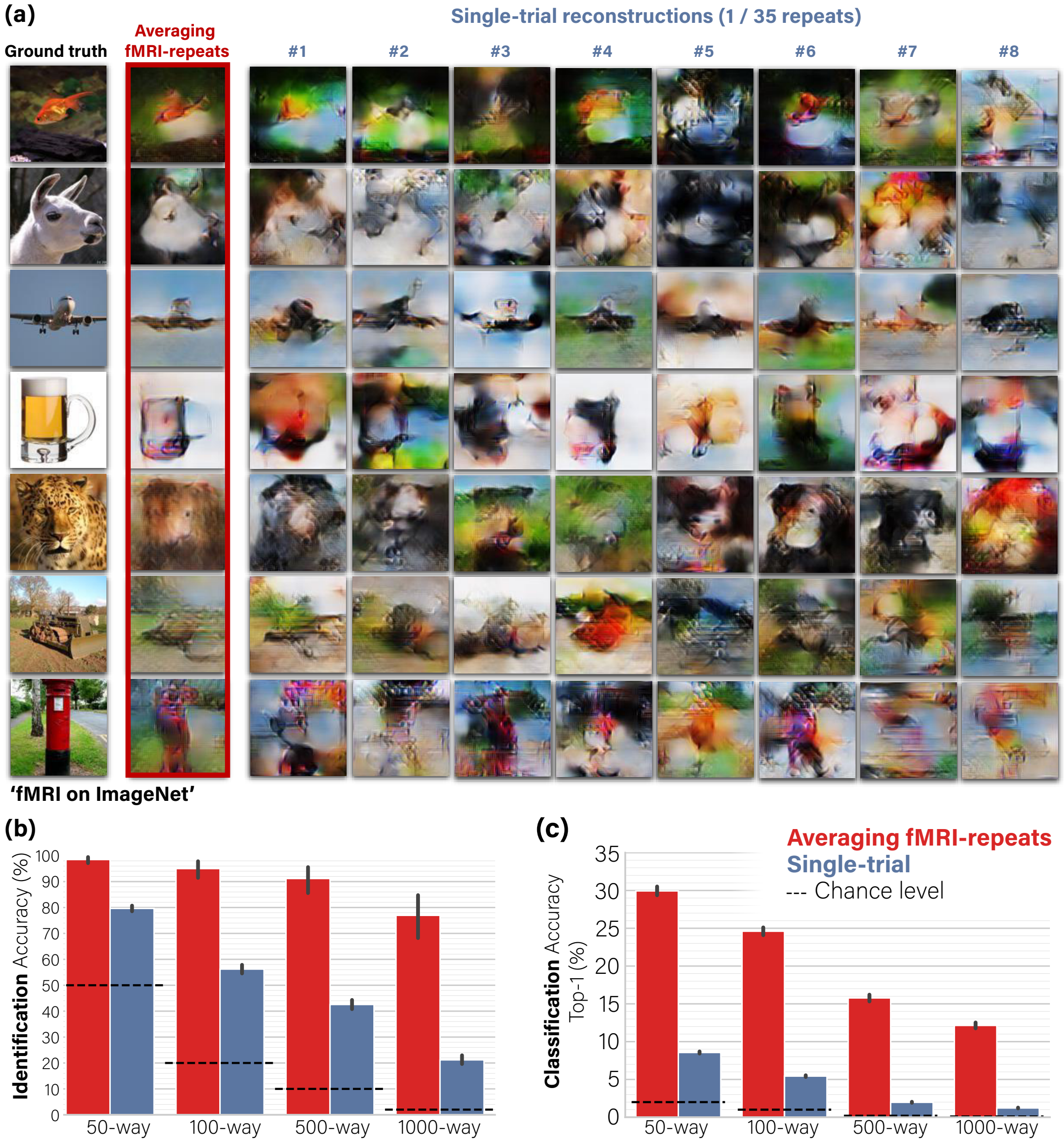


**Figure 10. Single-trial reconstruction, identification, and classification.** *(a) Reconstructed images from single-trial fMRI contrasted with those inferred from repeat-averaged fMRI data (showing eight random single-trials out of 35). The single-repeat-based reconstructions are significantly degraded and highly variant across repeats.* ***(b)*** *Identification results for all the single-trial-based reconstructions (50x35) compared with the average-based ones.* ***(c)*** *Similar to* ***(b)*** *but for semantic classification. Results are demonstrated for 'fMRI on ImageNet' [1] Subject 3 (with the highest noise ceiling).*

### Voxel selection

To account for the differences in voxel reliability, we considered two measures. The first measure was voxel signal to noise ratio (SNR). By 'signal' we refer to the response variance when different images were presented, while 'noise' is defined as the variance across trials where the same image was presented. The second measure was reproducibility across repeated test trials (cf. intra-subject reproducibility in Wen et al. [3]). This refers to correlation of voxel response in a specific trial with the average response across trials; each series consists of 50 responses representing the number of distinct test images. We found full agreement between these two measures.

We accounted for voxel reliability distribution in our Decoder model by encouraging strong sparsity on the weights which are associated with low-SNR voxels. We used the inverse-SNR of each voxel to weigh the lasso regularization on the weights which are associated with it (weights from the fully-connected layer).

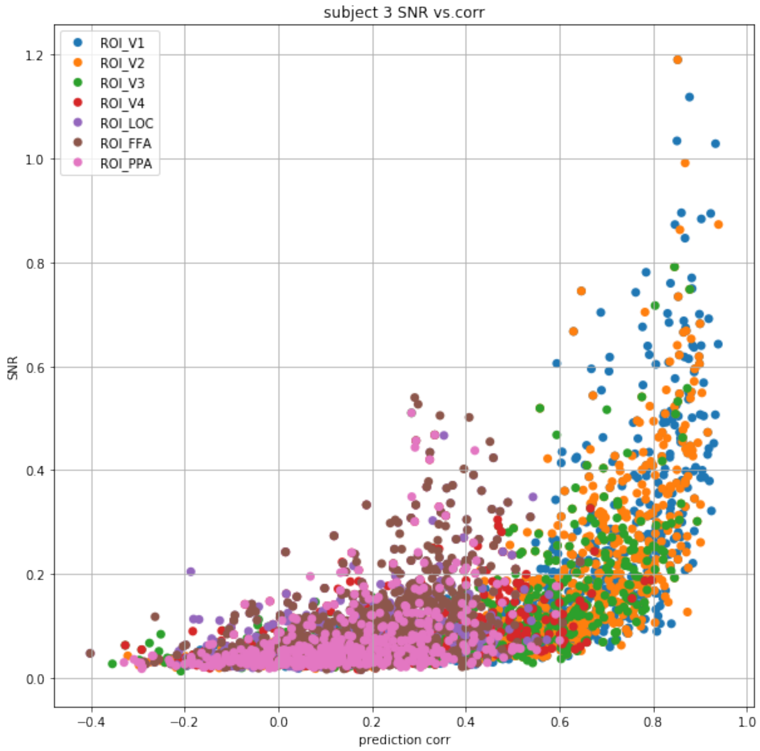


**Figure 11. Earlier visual areas have higher encoding prediction accuracy and higher SNR.**

### Spatial Grouping

Our decoding model contains a fully connected layer that constitute the transition between vector form voxel representation and spatial feature maps tensor. Each voxel had $\left\{w_{ij}^c\right\}$ weights connecting it to feature map $c$ at location $i, j$. Using this notation we define the spatial grouping loss per voxel,

$$\mathcal{L}_{SG} = \sum_{ij} \sqrt{\sum_{c} \kappa_{ij}^c} \tag{1}$$

$$\kappa_{ij}^c = (1-\alpha)\left(w_{ij}^c\right)^2 + \frac{\alpha}{4} \sum_{\Delta_i, \Delta_j \in \{-1,1\}} \left(w_{i+\Delta_i, j+\Delta_j}^c\right)^2, \tag{2}$$

where we used $\alpha = .5$ to force the grouping. Equation 1 expresses $L1$ (sparsity) loss in the spatial dimensions and an $L2$ loss in the channel dimension. This encourages an all-or-none inclusion of feature maps across spatial locations depending on their selection. The grouping force is due to the second term in Equation 2. Intuitively, this term accepts the weights of the neighboring locations at a reduced cost.

Note that the above regularization is a particular case of group lasso solution as defined in [4].

where $\beta_j$ is a vector of the weights associated with a specific spatial location, denoted by $j$: these are the weights of that location and its four perpendicularly neighboring locations; $K_j$ is a diagonal positive definite matrix that we parameterize with $\alpha$.
